# Basis for the phototaxis sign reversal in the green alga *Chlamydomonas reinhardtii* studied by high-speed observation

**DOI:** 10.1101/2020.12.06.414052

**Authors:** Masako Nakajima, Kosuke Iizuka, Rina Takahashi, Noriko Ueki, Atsuko Isu, Kenjiro Yoshimura, Toshiyuki Nakagaki, Toru Hisabori, Katsuhiko Sato, Ken-ichi Wakabayashi

## Abstract

For organisms that respond to environmental stimuli using taxes, reversal of the tactic sign should be tightly regulated for survival. The biciliate green alga *Chlamydomonas reinhardtii* is an excellent model for studying reversal between positive and negative phototaxis. *C. reinhardtii* cells change swimming direction by modulating the balance of beating forces between their two cilia after photoreception at the eyespot; however, it remains unknown how they reverse phototactic sign. In this study, we observed cells undergoing phototactic turns with a high-speed camera and found that two key factors determine the phototactic sign: which of the two cilia beats more strongly for phototactic turning and when the strong beating starts. The timing of the strong ciliary beating is suggested to be regulated by ROS-regulated switching between the light-on and light-off responses at the eyespot, which leads to the switching between positive and negative phototaxis. This idea is supported by a mathematical model that introduces the timing of the strong ciliary beating after photoreception.

## INTRODUCTION

The biciliate unicellular green alga *Chlamydomonas reinhardtii* is an excellent model organism to study how organisms respond to changing light conditions. *C. reinhardtii* shows a distinct light-induced behavior known as phototaxis, in which cells swim either toward or away from a light source (called positive or negative phototaxis, respectively). The direction of phototaxis, referred to here as the sign (either positive or negative), can be reversed. The regulation of this sign reversal is thought to be important for the viability of photosynthetic algae, but its mechanism is not well understood.

In *C. reinhardtii*, the light signal for phototaxis is received by the eyespot (Foster & Smyth, 1980), an organelle that appears under a microscope as an orange spot near the cell equator. It consists of carotenoid-rich granule layers, and a small area of plasma membrane in which channelrhodopsin (ChR) molecules are localized (Fig. 1A). The carotenoid-rich granule layers (CLs) function as a quarter-wave plate that reflects light (Foster & Smyth, 1980; Ueki et al., 2016), while ChR molecules are light-gated ion channels (Nagel et al., 2002; Nagel et al., 2003; Sineshchekov, Jung, & Spudich, 2002; Suzuki et al., 2003). Because of the relative position of these two components, the eyespot perceives light with high directionality. When a light signal arrives from the ChR-facing side, the light signal is amplified by reflection from the CLs; conversely, light signals are blocked by the CLs when coming from the CL-facing direction. In addition, while swimming, the cell rotates around its anterior-posterior axis. Directional photoreception by cells that swim with bodily rotation enables them to accurately detect and move in the direction of light stimuli.

**Fig. 1.**
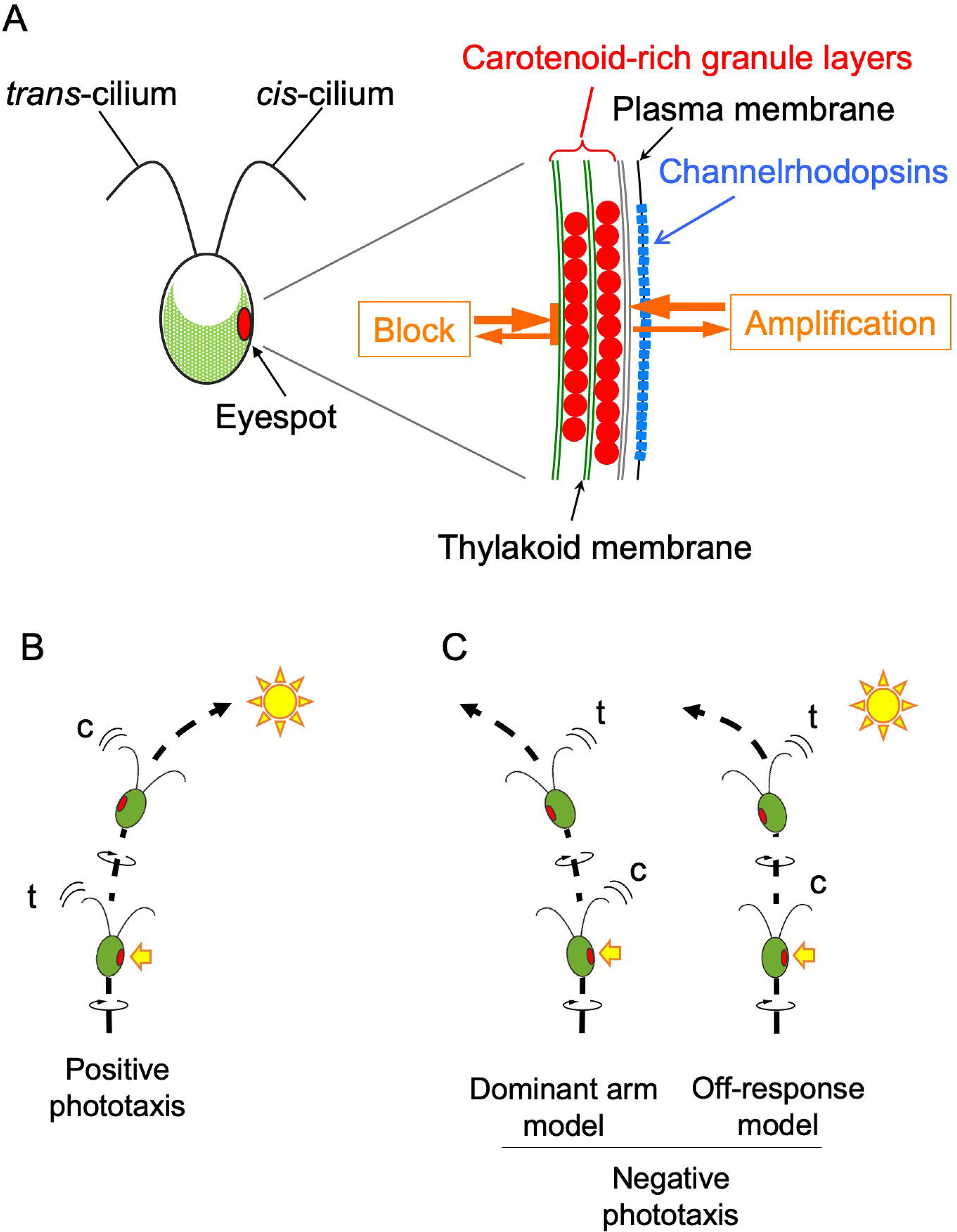
Schematic images of a *Chlamydomonas reinhardtii* cell and its phototaxis. **(A)** Schematic image of *C. reinhardtii* cell and the eyespot. The cilium closest to the eyespot is called *cis*-cilium, whereas the one farthest from the eyespot is called *trans*-cilium. The eyespot is composed of the carotenoid-rich granule layers that function as a light reflector, and channelrhodopsin molecules aligned in the plasma membrane. Channelrhodopsin functions as a light-gated cation channel. The light signal coming from the outside of the cell is reflected by the carotenoid granule layers and amplified, whereas that coming through the cell is blocked. (B) The current model to explain positive phototaxis. After photoreception at the eyespot (yellow arrow), [Ca^2+^]_i_ in the cilia increases and the *trans*-cilium (t) becomes dominant. After a half rotation of the cell around its central axis, the light signal to the eyespot is blocked, [Ca^2+^]_i_ in the cilia decreases, and the *cis*-cilium (c) becomes dominant. (C) Hypothetical models for negative phototaxis. In the “Dominant arm model”, Ca^2+^-sensitivities of two cilia are assumed to be reversed from the positively phototactic cell shown in (B); the *cis*-cilium becomes dominant after photoreception. In the “Off-response model”, after photoreception, the *trans*-cilium becomes dominant when the eyespot faces opposite to the light source.

Previous studies suggest that the phototaxis pathway in *C. reinhardtii* consists of three steps: 1) photoreception by ChRs, 2) an increase in intraciliary [Ca^2+^], and 3) a change in the balance of beating between the two cilia (a.k.a. flagella). After photoreception, the forces generated by the two cilia of *C. reinhardtii* become imbalanced, changing the cell’s swimming direction. These two cilia can be distinguished by their position relative to the eyespot, with the one nearest the eyespot called the *cis*-cilium, and the other called the *trans-cilium*. A currently prevailing model explains the mechanism underlying positive phototaxis as follows: When the eyespot faces a light source during swimming with bodily rotation, ChRs open to allow for Ca^2+^ entry, intraciliary [Ca^2+^] increases, and the *trans*-cilium starts to beat more strongly than the *cis*-cilium, by increasing beating amplitude and/or frequency (Kamiya & Witman, 1984; Rüffer & Nultsch, 1991). The imbalance between the forces generated by the two cilia tilts the cell’s swimming direction toward the eyespot-bearing side (i.e., in the direction of the light source). After 180° of rotation, the eyespot stops receiving light due to the shielding of CLs, the ChRs close, intraciliary [Ca^2+^] decreases due to some ion pumping activity, and the force generated by the *cis*-cilium increases while that of the *trans*-cilium decreases, resulting in the cell’s swimming direction tilting toward the light-source. By repeating this process, the cell will swim toward the light source, displaying positive phototaxis (Fig. 1B). However, in contrast to this widespread model, phototactic turns may also be initiated when the eyespot faces away from the light source (i.e., when the eyespot is shaded) (Isogai, Kamiya, & Yoshimura, 2000).

How, then, can the sign of phototaxis be reversed? Two possibilities exist: (i) The *cis*-cilium, rather than the *trans*-cilium, beats more strongly when [Ca^2+^]_i_ increases upon photoreception (referred to as “the dominant arm model”); and (ii) after photoreception, the *trans*-cilium beats more strongly than the *cis*-cilium when the cell makes a half bodily rotation and the eyespot faces the opposite side to the light source (referred to as “the off-response model). These possibilities should be verified by observing cell behavior after inducing positive or negative phototaxis.

Several factors are known to induce sign-switching of phototaxis in *C. reinhardtii*, including light intensity (Feinleib & Curry, 1971), extracellular Ca^2+^ concentration (Morel-Laurens, 1987), circadian rhythms (Kondo, Johnson, & Hastings, 1991), photosynthetic activities (Takahashi & Watanabe, 1993). Of these factors, the amount of cellular reactive oxygen species (ROS) changes the sign of phototaxis most intensely (Wakabayashi et al., 2011). Applying membrane-permeable ROS reagents or membrane-permeable ROS scavengers to cells can strongly bias the phototactic sign to positive or negative, respectively.

In this study, to test the two possibilities (i) and (ii) above, we carried out high-speed observations of cells turning during positive or negative phototaxis, after treatment with either a membrane-permeable ROS or a membrane-permeable ROS scavenger. Our results showed that the both (i) and (ii) are the cases. The innate phototactic sign of a strain can be explained by the case (i): after photoreception, we observed that positively phototactic strain cells beat the *trans-cilium* more strongly, whereas negatively phototactic strain cells beat the *cis*-cilium more strongly. When the same strain reverses the phototactic signs, the case (ii) applies: for example, the positively phototactic strain cells showed negative phototaxis by beating the trans-cilium more strongly when the eyespot faced opposite to the light source. The case (ii) can be interpreted in two ways: (a) the strong beating was induced by a light-off response of the eyespot, or (b) the light-on response of the eyespot was delayed by the time it takes for the cell to make a half rotation. We developed a mathematical model that satisfies the cases (i) and (ii). The model strongly suggested that the case (a) would be the case, which was confirmed by additional experiments using the slow-swimming cells.

## RESULTS

### Two hypotheses to explain the sign change of phototaxis

The mechanism underlying positive phototaxis can be explained as described in the Introduction (shown in Fig. 1B). To elucidate the mechanism of negative phototaxis, we considered two models (Fig. 1C), in which the dominant cilium is a key factor (defined here as the cilium that begins to beat more strongly after photoreception than the other). In the first model (the dominant-arm model), we assume that the relationship between the dominant cilium and [Ca^2+^]_i_ is reversed, such that the *cis*-cilium, rather than the *trans*-cilium, becomes dominant when [Ca^2+^]_i_ is increased. If the *cis*-cilium beats more strongly than the *trans*-cilium after light perception (which elicits Ca^2+^ influx), the cell will undergo negative phototaxis. In the second model (the off-response model), we assume that the relationship between the dominant cilium and [Ca^2+^]_i_ remains the same, but the onset of ciliary dominance (along with [Ca^2+^]_i_ increase) occurs when the eyespot faces away from the light source, and senses a light-off stimulus. In this case, the cell will also show negative phototaxis.

### Dominant cilium differs in strains with opposite phototactic signs

To determine which of these two models was more plausible, we observed the turning of *C. reinhardtii* cells using a high-speed camera (150 fps) linked to green light-emitting diode (LED) illumination sources at right angles. This system was an extension of the right-angle illumination system originally designed for the previous study (Isogai et al., 2000), which we here improved upon to observe the phototactic turnings and the eyespot position simultaneously by synchronizing the high-speed camera and the LED illumination. This allowed us to determine the exact time when a light stimulus was applied to a rotating cell, and the simultaneous orientation of the eyespot (visible as a bright spot under a dark-field microscope with an oil-immersion condenser). With this system, we first induced phototaxis in cells using the weaker stimulus of Light 1 (Fig. 2A). Then, the stronger Light 2 was illuminated at right angles to Light 1. When the eyespot in a swimming cell faced the light-source side of Light 2 (or the light side) after its illumination, we considered the cell to have perceived the light in that time frame (Fig. 2A). Then, from the position of the eyespot and the swimming path, we assessed which cilium beat more strongly (SI Movies S1-S4).

**Fig. 2.**
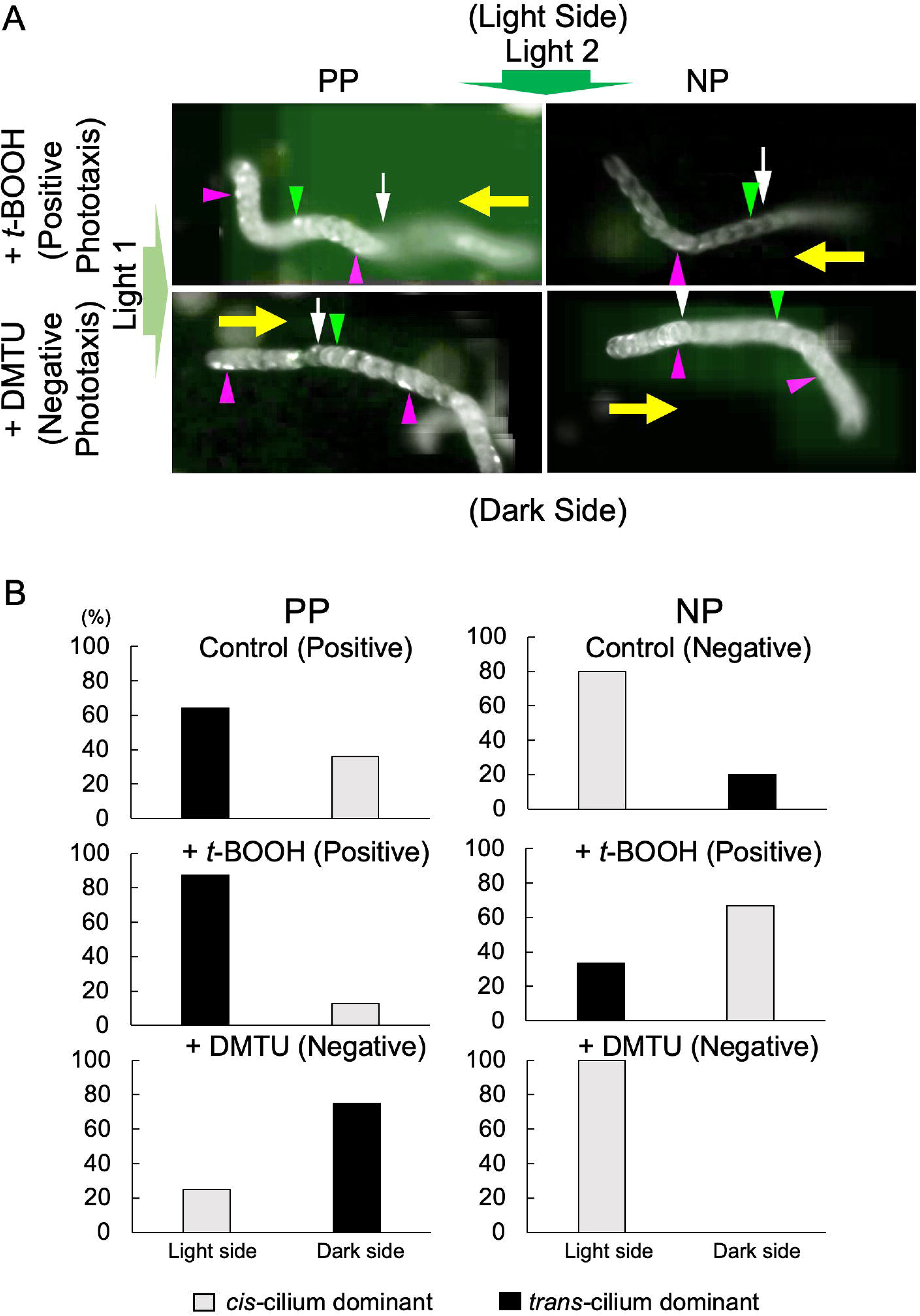
Eyespot position relative to the cell trajectory during phototaxis. **(A)** Phototactic turnings of PP and NP cells after treatment with t-BOOH or DMTU. Images of a swimming cell, taken at 150 fps, are superimposed every 0.13 sec. First, Light 1 (weak green light from the left) was illuminated to induce positive (top; swimming to the left) or negative (bottom; swimming to the right) phototaxis. After Light 2 (strong green light from the top) was turned on, the cell changed its swimming direction. The timing of the onset of Light 2 was at the white arrows/arrowhead. The eyespot facing Light 2 is shown with green arrowheads, whereas that facing opposite to Light 2 is shown with magenta arrowheads. The former case is classified as “the light side”) and the latter “the dark side” in **(B).** Yellow arrows show the swimming directions. (The cells whose trajectories intersect with the superimposed cells were removed from the image in the process creating the superimpose. See SI Movies S1-4 for the raw data.) **(B)** The proportion of the dominant cilium and the side where the onset of the ciliary dominance occurred. PP cells showing positive phototaxis (control and +*t*-BOOH) or negative phototaxis (+DMTU) and NP cells showing negative phototaxis (control and +DMTU) or positive phototaxis (+*t*-BOOH) were observed (n=8~28 per condition).

We observed the eyespot position during phototactic turning in two genetically different strains, CC-124 (a <u>n</u>egatively phototactic strain, here referred to as NP) and CC-125 (<u>p</u>ositively phototactic strain, PP) (see Materials and Methods). We analyzed only those cells that showed phototactic turnings of one full self-rotation after photoreception. We found that PP cells usually showed positive phototaxis. Similar to the PP cell after treatment with *t*-BOOH that induces positive phototaxis in Fig. 2A, a typical control PP cell changed its swimming direction immediately after it detected Light 2 (green arrow position). When turning toward the direction of the light source, more than half of the cells performed phototactic turning while the eyespot was located on the light side (i.e., the *trans*-cilium became dominant immediately after photoreception) (Fig. 2B).

In contrast, NP cells usually showed negative phototaxis. Similar to the NP cell after treatment with DMTU that induces negative phototaxis in Fig. 2A, a typical control NP cell changed its swimming direction immediately after it detected Light 2, similar to the PP cell. When turning against the direction of the light source, most cells performed turns while the eyespot was located on the light side, i.e. the *cis*-cilium became dominant immediately after photoreception (Fig. 2B). These observations suggest that the difference between PP and NP cells can be explained by the “Dominant arm model.” The cilium that becomes dominant after photoreception is genetically determined. Therefore, if the dominant cilium is the *trans*-cilium, the strain tends to show positive phototaxis, while if it is the *cis*-cilium in another strain, it tends to show negative phototaxis.

### Reversal of phototactic sign by “off-response” of the dominant cilium

Next, we examined the effect of reagents that change the cellular ROS, affecting the phototactic sign. After treatment with 0.2 mM *t*-BOOH, a membrane-permeable ROS reagent that induces positive phototaxis in both strains, most PP cells made phototactic turns while the eyespot was located on the light side, while most NP cells made phototactic turns while the eyespot was located on the dark side (i.e., the side opposite the light-source) (Fig. 2B) (SI Movie S1, S2). Fig. 2A shows the representative data, and the *t*-BOOH-treated NP cell changed its swimming direction after the cell made a half-self-rotation (magenta arrow). In contrast, after treatment with 75 mM DMTU, a membrane-permeable ROS scavenger that induces negative phototaxis in both strains, most PP cells made phototactic turns while the eyespot was located on the dark side, and most NP cells made phototactic turns while the eyespot was located on the light side (SI Movie S3, S4). In Fig. 2A, the DMTU-treated PP cell changed its swimming direction after the cell made a half-self-rotation (magenta arrow).

Thus, the sign-reversal of phototaxis in both strains can be explained by the off-response model; i.e., *t*-BOOH induces the onset of *cis*-cilium dominance in the dark side in NP cells and DMTU induces that of *trans*-cilium dominance in the light side in PP cells. Simply put, the reversal of genetically determined phototactic signs in a given strain is achieved by reversing the eyespot position relative to the light source (the light or dark side) when its genetically determined dominant cilium starts to show stronger beating than the other after light stimulation. We found that the dominant cilium after photoreception of PP cells was always the *trans*-cilium, while that of NP was always the *cis*-cilium. In each strain, the sign-switching of phototaxis was caused by the onset of strong beating of the dominant cilium on the dark side.

Therefore, these phototactic turning events can be categorized into two cases for each sign of phototaxis. For cells displaying positive phototaxis: in (Positive Case 1), phototactic turning occurred while the eyespot was located on the light side. In this case, the *trans*-cilium must have become dominant in response to the light-on stimulus (such as PP cells without the ROS-reagents or with *t*-BOOH). In (Positive Case 2), phototactic turning occurred while the eyespot was located on the dark side. In this case, the *cis*-cilium must have become dominant in response to the light-off stimulus (such as NP cells with *t*-BOOH). For cells displaying negative phototaxis: in (Negative Case 1), phototactic turning occurred while the eyespot was located on the light side. In this case, the *cis*-cilium must have become dominant in response to the light-on stimulus (such as NP cells without the ROS-reagents or with DMTU). In (Negative Case 2), phototactic turning occurred while the eyespot was located on the dark side. In this case, the *trans*-cilium must have become dominant in response to the light-off stimulus (such as PP cells with DMTU).

### Eyespot position in the helical paths

Regarding changes in the balance of beating between the two cilia during phototactic turning, we first wanted to examine the force balance in unstimulated cells. Under homogeneous light conditions, *C. reinhardtii* cells swim in a helical path, due to the imbalanced force generation of the two cilia beating in slightly skewed planes (Fig. S1A). In previous work, Isogai et al. showed that the eyespot is located on the outer edge of the helical swimming paths in ~80 percent of positively phototactic cells (Isogai et al., 2000). In contrast, for negatively phototactic cells, ~50 percent of cells swim with the eyespot facing inside, whereas ~30 percent of cells swim with the eyespot facing outside (Isogai et al., 2000). These results suggest that eyespot position in the helical swimming path is partially correlated with the phototactic sign, but the correlation is not determinative. We thus investigated whether the eyespot position in the swimming paths is affected by ROS-modulating reagents. Most, but not all, cells swam with the eyespot outside the helix when they showed either positive phototaxis (Fig. S1B, Sign P). However, when cells showed negative phototaxis in the presence of DMTU, directing the eyespot inside the helix was observed almost as frequently as towards the outside (Fig. S1B, Sign N, +DMTU). Thus, we did not observe a strict correlation either between phototactic sign and eyespot position in the helical paths.

To elucidate if the dominant cilium after photoreception is also dominant in the swimming without light stimulation, we observed the PP and NP cells swimming under a microscope in the red background light. The results showed that ~60% cells swam with the eyespot facing outward of the helical path and the rest swam with it inward in both PP and NP cells (Fig. S1C). These data also suggest that the dominant cilium after photoreception is independent from that without photostimulation. These results are consistent with those of a mathematical model that describes the swimming behavior of *C. reinhardtii* given in the section of “Mathematical model to test the experimental data,” which shows that whether the eyespot faces outward or inward of the helical path does not affect the sign of phototaxis.

### Photoreceptor currents after treatment with the ROS-modulating reagents

A possible explanation for how ROS-modulating reagents induce the light-off response is that those reagents delay the response of the dominant cilium by slowing the opening of photo-gated channels (ChRs). If this delay is as long as the time required for a cell to perform a half rotation (~250 msec), then the phototactic sign of the cell would reverse. To test this possibility, we measured the photoreceptor current (PRC) in a population of *C. reinhardtii* cells by the method of Sineshchekov et al. (Sineshchekov et al., 1992; Sineshchekov et al., 1994) (Fig. S2A). Treatment with *t*-BOOH did not significantly affect the magnitude of PRC produced by a single flash of light stimulation, in either PP or NP cells (Fig. S2B, C). We also measured the time (delay) required for the generation of PRC after photostimulation, but detected no significant difference between the control and the *t*-BOOH-treated cells (Fig. S2B, C). In contrast, after treatment with DMTU, we found that PRC decreased, while delay time increased, in both strains (Fig. S2B, C). However, the increase in delay time was only ~1 msec. These results suggest that the light-off response is not caused by the delay in PRC generation.

### Mathematical model to test the experimental data

We have shown that there are two cases where the phototactic sign of *C. reinhardtii* is reversed: (i) dominant arm model case and (ii) off-response model case. In the off-response model case, however, there is another interpretation of the experimental results, namely, if *C. reinhardtii* has a time delay for the response to the light stimulus and the delay time is so long that the cell rotates half a turn about its anterior-posterior axis, the phototactic turning occurs when the eyespot faces on the dark region; we refer to this hypothesis as “the time delay model”. To examine whether the time delay changes the phototactic sign of the cell and determine whether the off-response model or the time delay model is more plausible for explaining phototactic turning when the eyespot facing on the dark side, we develop a mathematical model that express the time delay of the cell. Our modeling framework is based on previous models of phototaxis in *C. reinhardtii* and the multicellular green alga *Volvox carteri* (Bennett & Golestanian, 2015; Drescher, Goldstein, & Tuval, 2010, Leptos et al., 2018). We extend them by explicitly introducing a time delay into the ciliary response after a light stimulus (see SI Appendix A for details). Brief description of our mathematical model is below.

We approximate the cell as a rigid ball and define the body axes of the cell as in Fig. 3A, where **a, b** and **c** are unit vectors that are fixed to the body of the cell. We assume that a cell swims with cilia forward, with a constant speed *v*_0_ in the posterior direction **c**, i.e., **v** = *v*_0_**c**, and rotates with the angular velocity

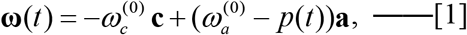

where 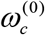 and 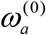 are constants, which represent the proper angular velocity of the cell under unstimulated conditions. The quantity *p* expresses the contribution from the change in the behaviors of the two cilia in response to the change in light intensity at the eyespot: when the *trans*-cilium beats stronger than the *cis*-cilium, *p* becomes positive, and vice versa. We assume that the response of dominant cilium to the light stimulus has some constant delay. Indeed, under the normal culture conditions the response of dominant cilium to the light has a short delay time around 30-40 msec (Rüffer & Nultsch, 1991; Witman, 1993). We extend this notion and allow the delay time *τ*_0_ to be any. We thus write the relationship between *p* and light intensity received by the eyespot, *I*, as

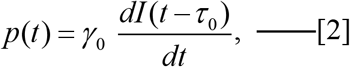

where *γ*_0_ is a constant that takes the value +1 or −1. *γ*_0_ = +1 means the *trans*-cilium is dominant (as in PP), while *γ*_0_ = −1 means the *cis*-cilium is dominant (as in NP). By changing the value of *g*, we express the dominant arm model. *τ*_0_ is the delay time of the onset of the ciliary dominance after photoreception. By changing the value of *τ*_0_ we can investigate the effects of the time delay of dominant cilium to light stimulus on phototaxis. In the mathematical description, the light-off response model in fact coincides with the dominant arm model, because the onset of dominant cilium at step-down stimulus (*γ*_0_ = +1 and *dI* / *dt* < 0) and the onset of exchanged dominant cilium at step-up stimulus (*γ*_0_ = −1 and *dI* / *dt* > 0) cannot be distinguished in this framework. Thus this model is used to compare the results of the time delay model and those of the other models, dominant arm model and off-response model.

**Fig. 3.**
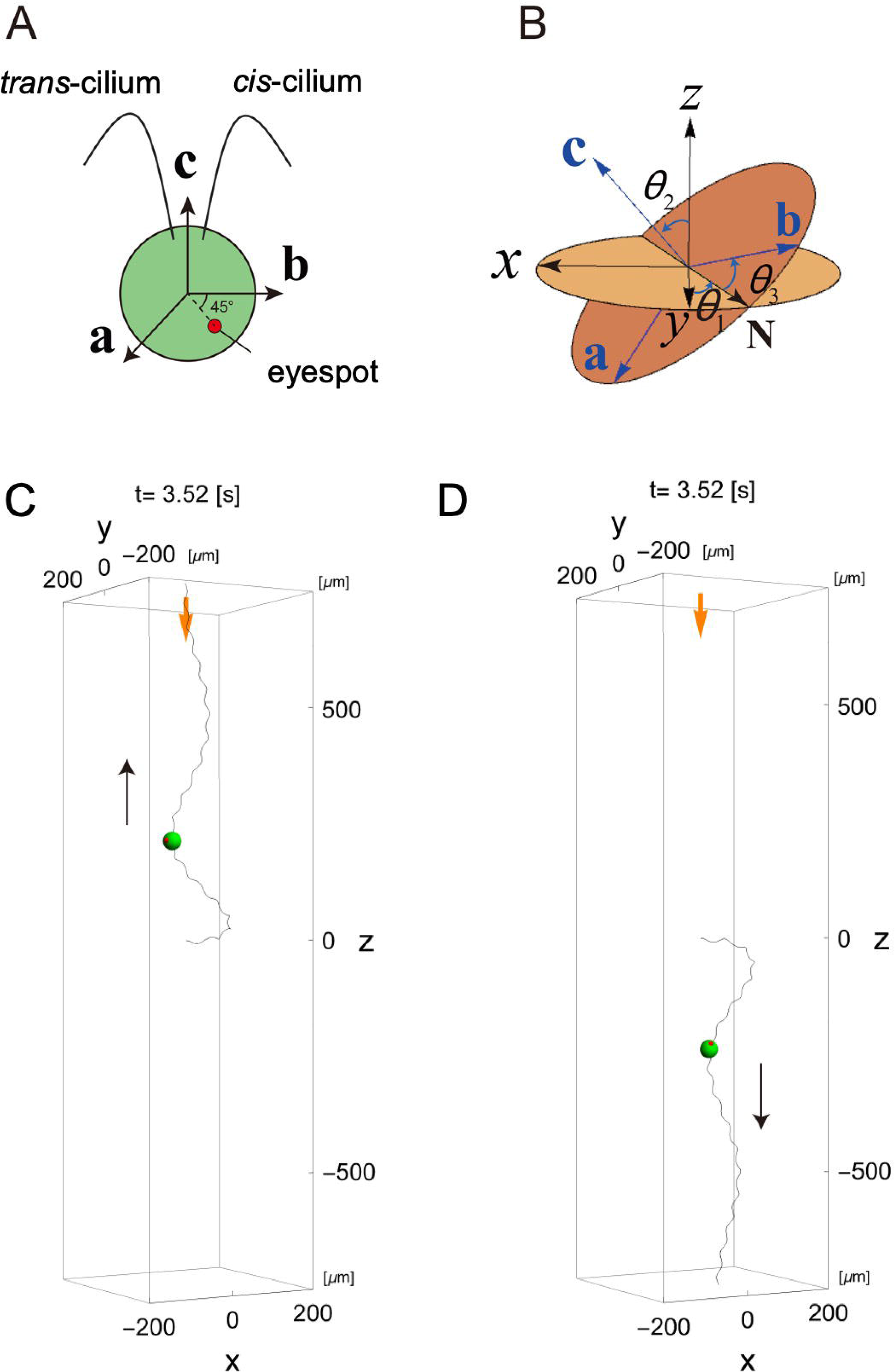
Mathematical model for *C. reinhardtii* phototaxis. **(A)** Definitions of the body axes of a *C. reinhardtii* cell. The vectors **a**, **b** and **c** are unit vectors that are fixed to the body of the cell. **a**, **b** and **c** are mutually orthogonal to each other, and **b** and **c** are within the ciliary beat plane. **b** is close to the side of *cis*-cilium. With them, the direction of eyespot is expressed as 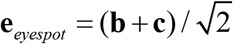. The directions of **a**, **b** and **c** evolve with time according to Eq. 1. **(B)** Definitions of the Euler angles that specify the directions of **a**, **b**, **c** for the x, y, z coordinate system that is fixed in space. *θ_1_* is the angle between the y-axis and the vector **N**, where **N** = **z**×**c**/|**z**×**c**|. *θ*_2_ and *θ*_3_ are angles between the z-axis and **c** and between **N** and **b**, respectively. When *θ*_1_ = *θ*_2_ = *θ*_3_ = 0, **a**, **b**, **c** axes coincide with *x*, *y*, *z* axes, respectively. **(C), (D)** Examples of initial trajectories of the cell obeying Eq. 1 (0 ≤ *t* ≤10), which indicate positive phototaxis (*τ*_o_ =0.08 sec, **(C)**) and negative phototaxis (*τ*_o_ =0.32 sec. **(D)**). The parameters are *γ*_0_ = 1 (the *trans*-cilium becomes dominant after photoreception at the eyespot), *I*_0_ = 0.5, *v*_0_ = 128 [μm/s], [1/s], *ω_c_* = −11.8 [1/s] and *ω_a_* = 5.8 [1/s]. The initial conditions are **r**(0) = (0,0,0), *θ*_1_(0) = 0, *θ*_2_(0) = −*π*/2 and *θ*_3_(0) = 0. Thick orange arrows show the direction of the light illumination, and thin black arrows show the swimming direction of the cell.

The light intensity *I* received by the eyespot is given by *I*(*t*) = *I*_0_ (−**e**_*light*_·**e**_*eyespot*_ +1) / 2, where *I*_0_ is the intensity of the light at the cell position. **e**_*light*_ and 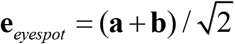 are unit vectors that represent the direction of the incident light and the eyespot, respectively. The time evolution equations for this model are closed in terms of the Euler angles *θ*_1_, *θ*_2_, *θ*_3_) that specify the directions of body axes, **a**, **b**, **c** (Fig. 3B) (Landau, 1976) and the position of the cell **r** = (*x, y, z*).

With this model we consider a simple situation where a parallel light comes from the positive z direction (Fig. 3B), which is expressed by **e**_*light*_ = (0,0,−1) and constant *I*_0_. If the cell goes in the positive z direction in the final state, this situation indicates positive phototaxis, while if the cell goes in the negative z direction in the final state, that is negative phototaxis.

This mathematical model in fact has two steady solutions (Eqs. S5 and S6 in Appendix B) for any parameters: one represents positive phototaxis, where the cell swims in the positive z direction with a constant speed, drawing a left-handed spiral trajectory. The other represents negative phototaxis, where the cell swims in the negative z direction with the same speed, drawing a left-handed spiral trajectory. Thus the model intrinsically involves both positive and negative phototaxis states. Depending on the values of model parameters, *γ*_0_, *I*_0_ and *τ*_0_, in the relation between *p* and *I*, the stability of the two steady solutions changes (Fig. S3), and one stationary solution is selected (Fig. 3C, D; SI Movies S5,S6). The stability of the steady solutions does not depend on *v*_0_, because the time evolution equations for (*θ*_1_, *θ*_2_, *θ*_3_) that spesify the direction of the cell are closed in terms of (*θ*_1_, *θ*_2_, *θ*_3_) only (see Eq. S4 in Appendix A) and do not contain *v*_0_.

Furthermore, from the steady solutions of this model we find the relationships between the parameters involved in our mathematical model (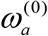, 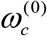, *v*_0_) and the quantities concerning the helical path of the cell observed in experiments as

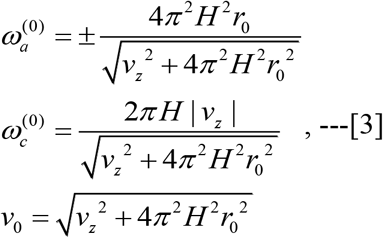

where *r*_0_, *H* and |*v_z_*| are the radius, Heltz, and speed of the cell along the axis of helical path drawn by the cell under homogeneous light conditions, respectively (see Appendix C for deitals). Using eq. 3 we can determine the values of model parameters 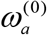, 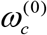, *v*_0_ from experimental data, except for the sign of 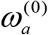. (Note that the first relation in Eq. 3 has plus minus sign.) The sign of 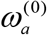 is in fact related to the relative position of the eyespot in the helical orbit of the cell (Appendix C), i.e., when 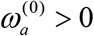 the eyespot face the outside of the helical orbit, while when 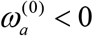 the eyespot faces the inside of helical orbit. This result holds independent of phototactic sign. That is, for each phototaxis, there are two cases where the eyespot faces the outside and inside of the helical path. In which direction the eyespot faces is determined by only the sign of 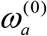. Using this property we can also determine the sign of 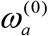 from the position of eyespot in the helical path. Most cells observed in our experiments exhibit that the relative position of eyespot to the helical orbit is the outside, so we take 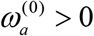 throughout this model. Hence the control parameters of this model turn out to be *γ*_0_, *I*_0_ and *τ*_0_.

To check whether this mathematical model exhibits definite phototaxis for given parameters, we numerically integrated the time evolution equations Eqs. S3 and S4 in Appendix A with some initial conditions with the Euler methd (Δ*t* = 1/10000). Figure 3 C,D give the trajectories of the cell described in our model for *τ*_0_ = 0.08 (C) and *τ*_0_ = 0.34 (D). We see that in Fig. 3C the cell goes in the positive z direction in the final state (positive pototaxis), while in Fig. 3D, the cell goes in the positive z direction (negative pototaxis). These final swimming directions do not depend on the initial conditions, i.e., even if the cell directed in any direction at the initial time, the final swimming direction (positive or negative phototaxis) is determined only by the values of the model parameters, *I*_0_, *γ*_0_ and *τ*_0_. To see how the swimming direction depends on the model parameters, we drew the phase diagram of the final velocity of the cell (Fig. 4), where the values of 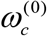, 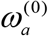 and *v*_0_ used here were obtained from experimental data on the PP cell using the relations Eq. 3. In Fig. 4 we see that when *γ*_0_ is changed from +1 to −1, the swimming direction of the cell becomes opposite. This means that when the dominant cilium is exchanged, the phototactic sign is reversed. This result is consistent with our experimental data in section _ and also with the existing works [interface, Bennett & Golestanian, 2015, PRL]. Whenever the sign of *γ*_0_ is changed, the phototactic sign is also changed, which comes from the symmetry of the mathematical model.

**Fig. 4.**
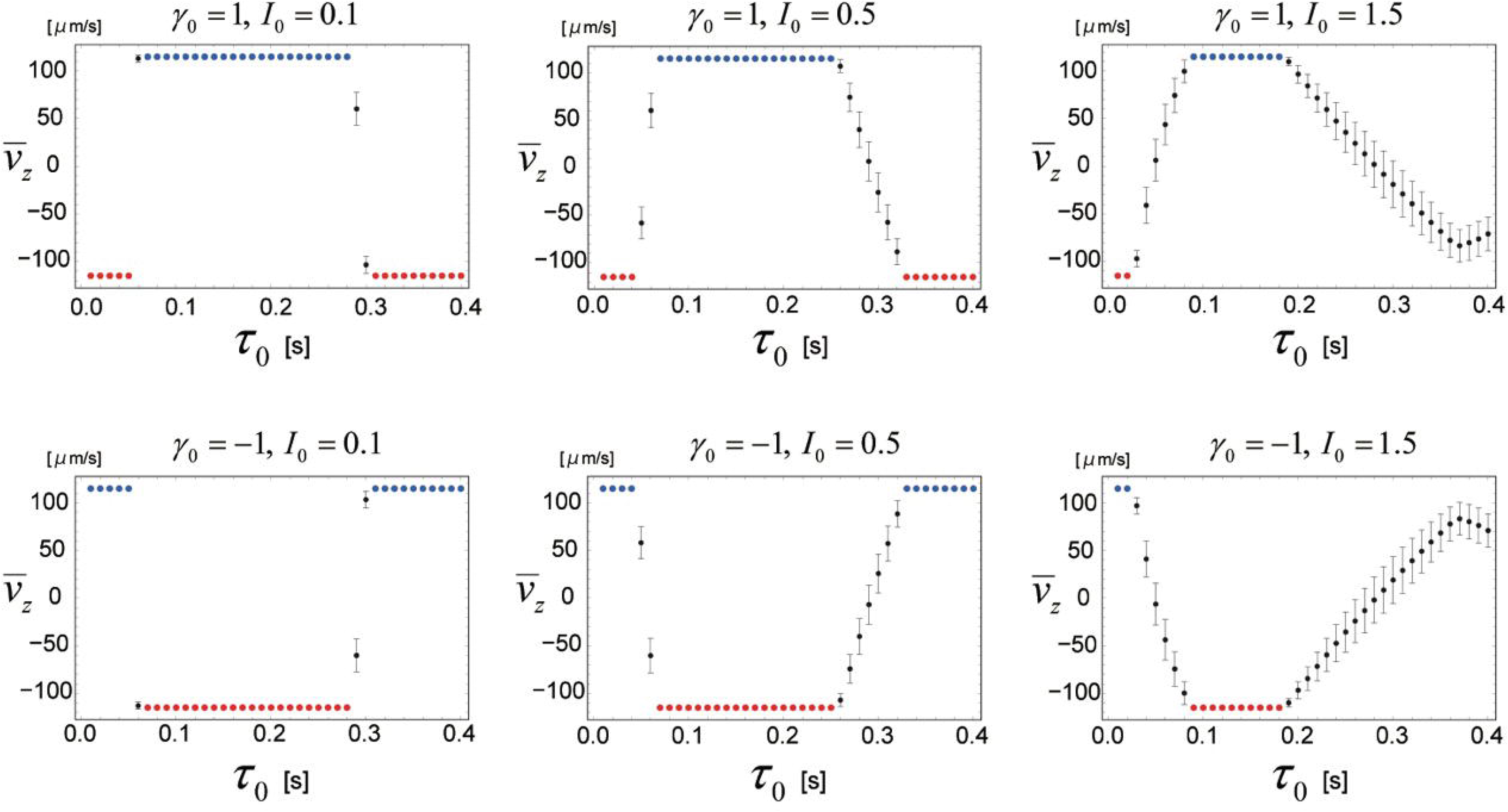
Sign-switching of phototaxis in the mathematical model. The mean velocity 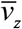 of the steady-state of the cell after a long-time simulation as a function of the delay time *τ*_0_ for various values of *γ*_0_ and *I*_0_. For each set of system parameters, only one steady state of Eq. 1 realizes, which does not depend on the initial conditions of *θ*_1_, *θ*_2_, *θ*_3_, **r**. 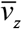 changes with *τ*_0_; especially, the sign of 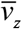 (=the sign of phototaxis) changes with *γ*_0_ and *τ*_0_. Blue dots are the states where solution S5 realizes, while red dots are the states where solution S6 realizes. Black dots with bars are the states where solutions other than Eqs. S5 or S6 are achieved, in which *v*_z_(*t*) oscillates in time. The bars indicate the standard deviation of *v_z_* of the steady-state. The parameter values used are *v*_0_ = 128 [μm/s], [1/s], *ω_c_* = −11.8 [1/s] and *ω_a_* = 5.8 [1/s]. The initial conditions are: **r**(0) = (0,0,0), *θ_i_*(0) = *δ_i_* with random numbers *δ_i_* ∈ [0,2*π*] for *i* = 1,2,3, and *dI*(*t*) / *dt* = 0 for 0 ≤*t*≤*τ_0_*. For the discretization of Eq. S1, the Euler method was used (Δ*t* = 1/10000). The model suggests that, when *γ*_0_ =1 (i.e. the *trans-cilium* is dominant), the cell shows positive phototaxis (i.e. *γ*_0_ is positive) when the dominant cilium beats stronger than the *cis*-cilium with the delay (*τ*_0_) 60~280 ms. Similarly, when *γ*_0_=-1 (i.e. the *cis*-cilium is dominant), the cell shows negative phototaxis under the same conditions.

When we fix the values of *I*_0_ and *γ*_0_ and change only *τ*_0_, the final swimming direction also changes (Fig. 4). For example, let us see the case of *I*_0_ = 0.1 and *γ*_0_ = +1 in Fig. 4. In this case photatctic sign becomes plus when 0.06 <*τ*_0_<0.3 [sec], and the phototactic sign becomes minus when 0.3 <*τ*_0_<0.4 [sec]. Since the mean self-rotation frequency of the PP cell about the anterior-posterior (AP) axis in experiments is about 2 Hz (see Table 1), the time needed for the cell to rotate about the AP axis one-half is about 0.25 [sec]. Thus, the above results of numerical simulations are basically consistent with the picture of the time delay model, i.e., we can consider that for 0<*τ*_0_<0.25 the eyespot faces on the bright side, while for 0.25 <*τ*_0_<0.5 the eyespot faces on the dark side. That is, there is a possibility that the behaviors of the cells obaserved in experiments given in section may be the result of delayed response of the cell to the light stimulus.

**Table 1.** Ciliary beating frequencies and bodily rotation cycles of strains used.

|  | PP | NP | *ida4* | *oda1* |
| --- | --- | --- | --- | --- |
| Ciliary Beating Frequency (Hz)<br>(Mean ± SEM, n=3 measurements) | 55.0 ± 0.6 | 55.5 ± 1.0 | 51.6 ± 2.6 | 24.3 ± 0.6 |
| Bodily rotation cycle (Hz)<br>(Mean ± SD, n=10 cells) | 2.11 ± 0.27 | 1.89 ± 0.27 | 1.25 ± 0.22 | 0.72 ± 0.09 |

### Off-response or delayed response?

In the Off-response model, ciliary dominance starts to occur on the dark side. We interpreted this as an immediate response to a light-off signal. However, it may also be a delayed response to a light-on signal. If the delay time *τ*_0_ is fixed between 0.25 sec and 0.5 sec, a delayed response to a light-on stimulus would just look like a light-off response without a delay. To assess which of these two interpretations is more plausible, we conducted two experiments and a simulation.

First, we observed phototaxis in slow-swimming mutants *ida4* (lacking inner-arm dynein subspecies a, c and d) and *oda1* (lacking entire outer-arm dynein) after treatment with ROS-modulating reagents (Kamiya, 1988; Kamiya et al., 1991; Takada et al., 2002). As the bodily rotation of the cell is caused by the slightly three-dimensional beating of the two cilia, slow-swimming mutants show bodily rotation with a longer rotation cycle time, as we observed (Table 1). If the delay time is between 0.25 and 0.50 sec, those slow-swimming mutants may differ in phototactic sign from the wild-type PP and NP strains. We found that both *oda1* and *ida4* cells tended to display positive phototaxis under neutral or oxidizing conditions, and negative phototaxis under reducing conditions, similarly to the PP strain of the wild type (Fig. 5A, B). However, the sign-reversal in the slow-swimming mutants was not as clear as in the wild type, especially after treatment with the ROS-scavenger. It is possible that the force generation for steering is weaker under these conditions in the slow-swimming mutants.

**Fig. 5.**
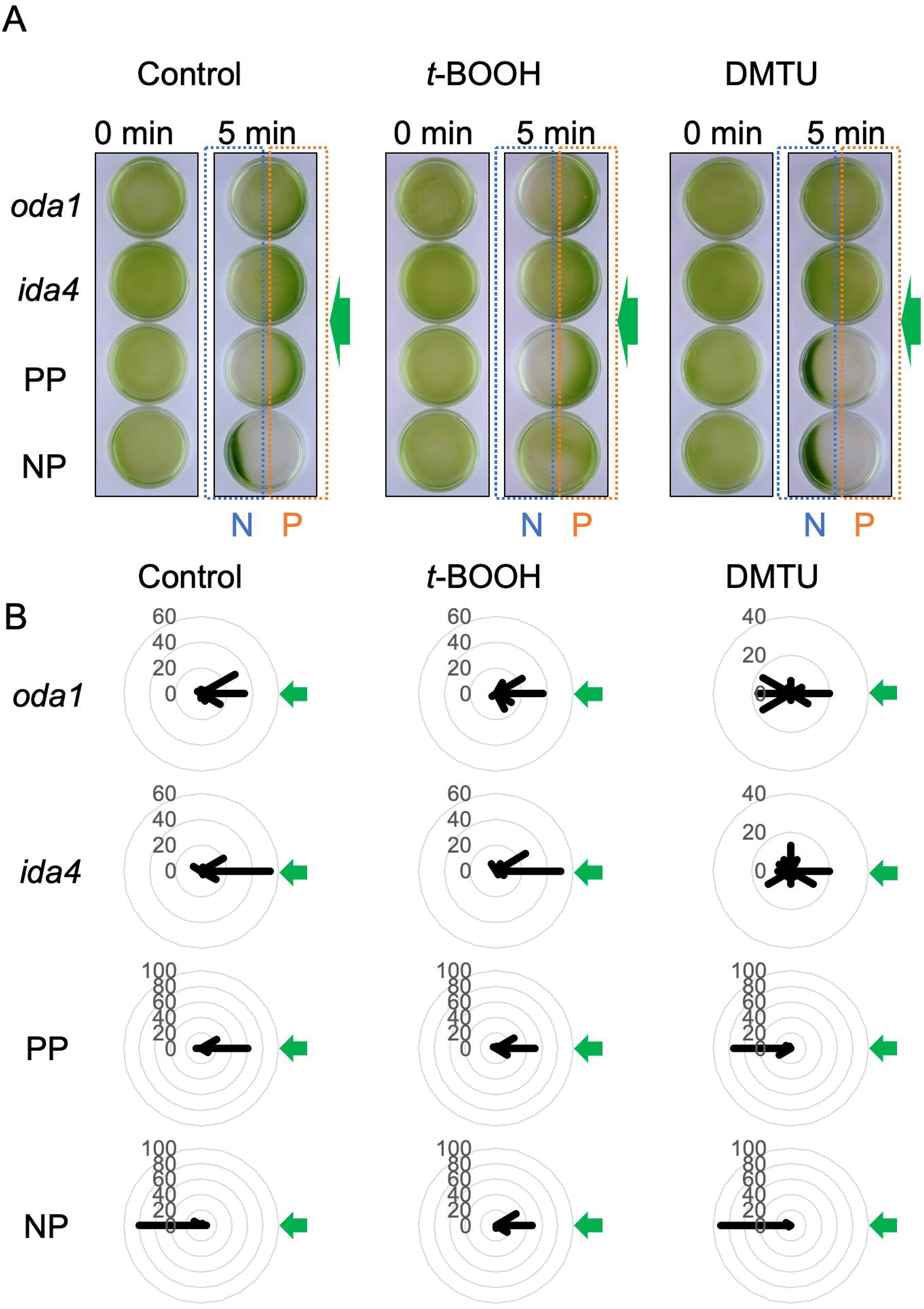
Phototaxis assay of the slow-swimming cells. **(A)** PP, NP, *oda1*, and *ida4* cell suspensions put in Petri dishes with or without ROS-modulating reagents (0.2 mM *t*-BOOH or 75 mM DMTU) were illuminated by green LED (λ=525 nm, 30 μmol photons m^-2^ s^-1^) from the right (green arrows) for 5 min from the right. Cells showing positive phototaxis are accumulated in the right halves of the dishes (orange boxes with “P”) and those showing negative phototaxis are accumulated in the left halves of the dishes (blue boxes with “N”). **(B)** Polar histograms depicting the percentage of cells moving in a particular direction relative to light illuminated from the right (green arrows), with or without treatment with ROS-modulating reagents (12 bins of 30°; n = 30 cells per condition).

Thus, we also observed phototaxis of PP cells after treatment with DMTU in a viscous medium. If the negative phototaxis of PP cells is achieved by the delayed response, the DMTU-treated PP cells may not be able to show negative phototaxis when the self-rotation duration is prolonged. In the medium containing 4.5% or 9.0% Ficoll400, cells swam slowly (Table 2). Even in these media, the DMTU-treated PP cells showed clear negative phototaxis (Fig. 6). These results cannot be achieved by the delayed response.

**Table 2.** Swimming parameters of PP in a solution containing Ficoll400.

|  | PP in 4.5% Ficoll | PP in 9.0% Ficoll |
| --- | --- | --- |
| Ciliary Beating Frequency (Hz)<br>(Mean ± SEM, n=3 measurements) | 38.2 ± 0.21 | 26.8 ± 1.35 |
| Bodily rotation cycle (Hz)<br>(Mean ± SD, n=10 cells) | 2.13 ± 0.77 | 0.91 ± 0.24 |
| Swimming velocity ($\mu\text{m/s}$)<br>(Mean ± SD, n=20 cells) | 105.9 ± 31.9 | 50.7 ± 12.6 |

**Fig. 6.**
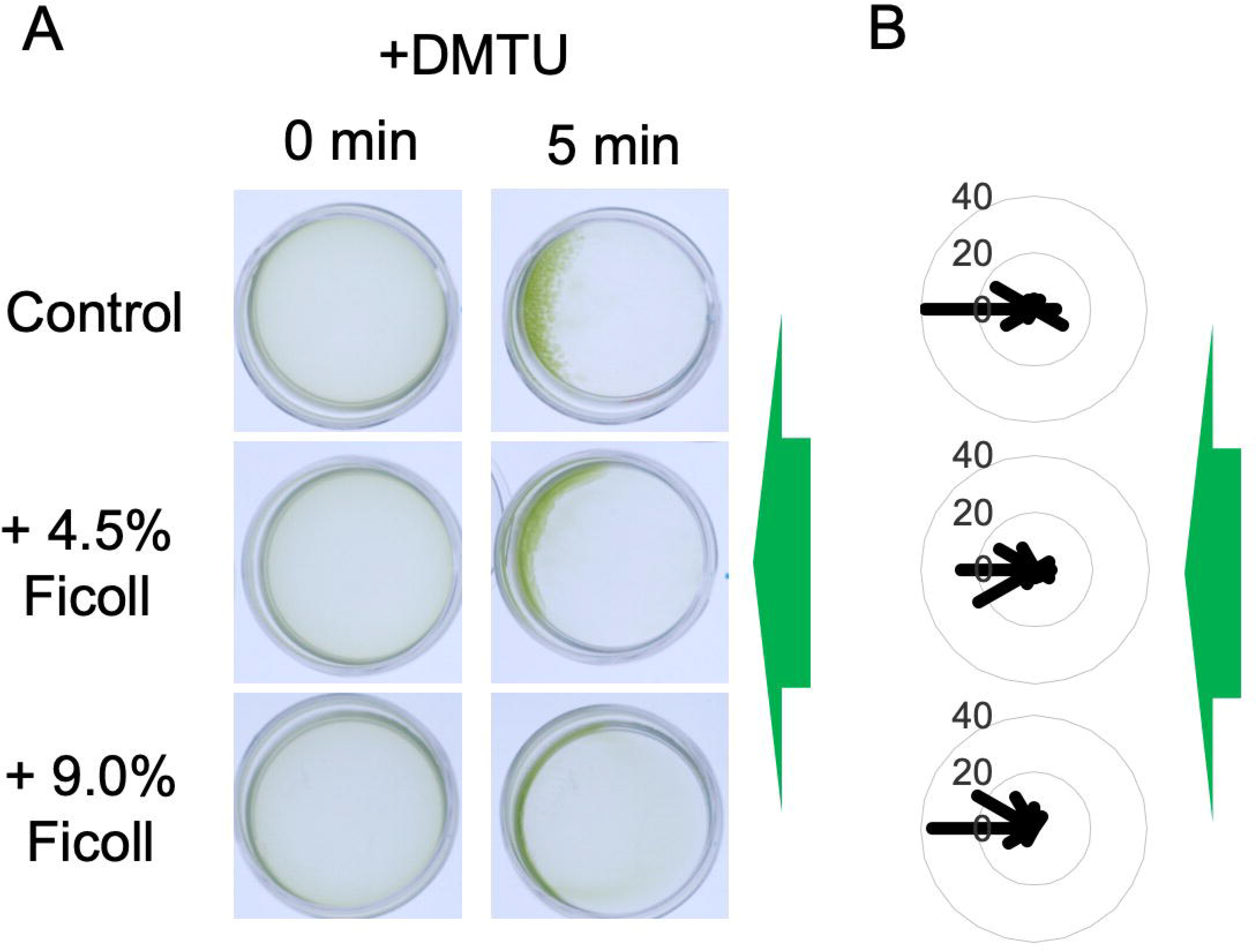
Phototaxis assay of PP cells in viscous media. (A) PP cells in viscous media (containing 4.5% or 9.0% Ficoll400) after treatment with 75 mM DMTU were illuminated by green LED (λ=525 nm, 30 μmol photons m^-2^ s^-1^) from the right (green arrows) for 5 min from the right. **(B)** Polar histograms depicting the percentage of cells moving in a particular direction relative to light illuminated from the right (green arrows) (12 bins of 30°; n = 30 cells per condition).

We also used our mathematical model to simulate the behavior of cells that self-rotate at a lower-than-normal frequency of 1.25 Hz and 0.72 Hz, frequencies of *ida4* and *oda1*, respectively (Table 1). Our simulation results indicated that when τ0 is between 0.075 and 0.5 sec (when self-rotation frequency is 1.25 Hz) or 0.25 and 0.5 sec (0.72 Hz), such cells do not change phototactic signs with a change in τ0, unlike the wild type cells that rotate at higher frequencies of ~2 Hz (Fig. 7). Furthermore, we carried out similar simulations using the self-rotation frequency of cells in viscous media (Table 2). In a medium containing 9.0% Ficoll400, the rotation frequency is ~0.91 Hz (Table 2). In this case, when τ0 is between 0.15 and 0.7 sec, cells do not change phototactic signs with a change in τ0 unlike cells in a regular medium (Fig. S4).

**Fig. 7.**
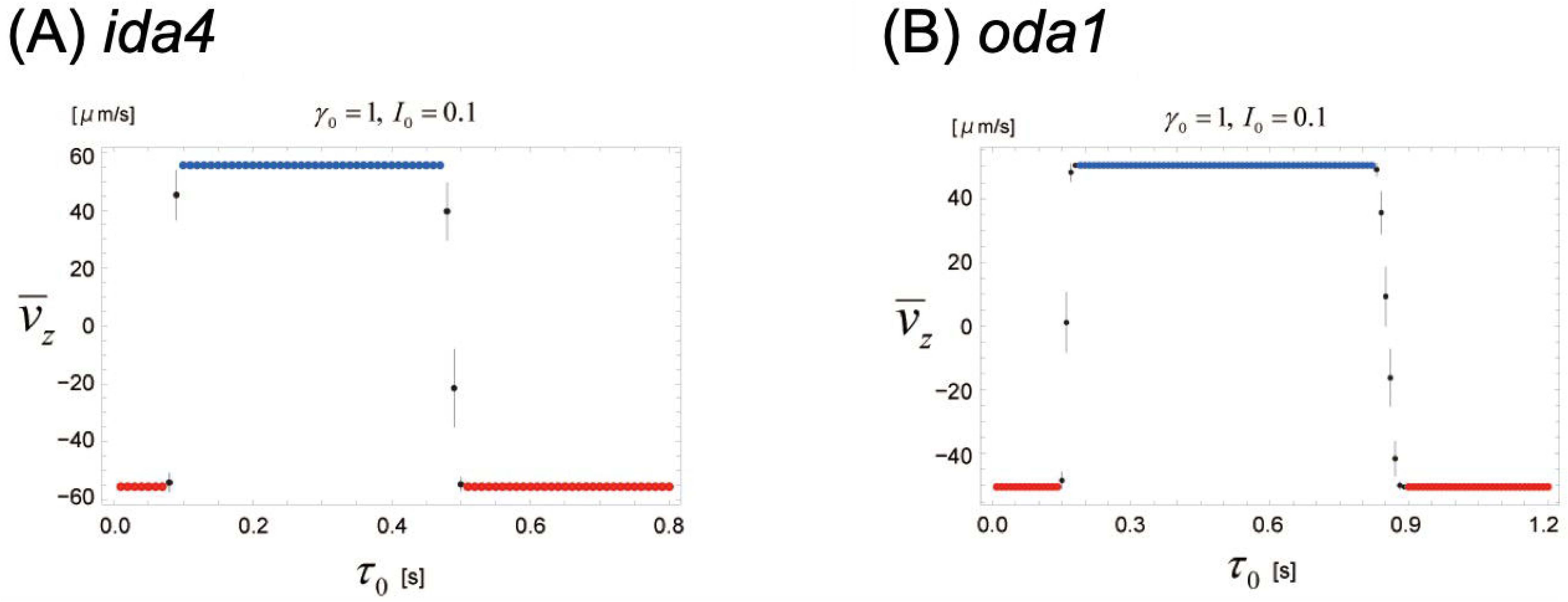
Sign-switching of phototaxis in a slow-swimming mutant in the mathematical model. The plots of the mean velocity 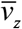 of the steady-state of the mathematical model (Eqs. S3 and S4 in SI Appendix A) as a function of the delay time *τ*_0_ for two cells: (A) *ida4* (rotation frequency 1.25 Hz) and (B) *oda1* (rotation frequency 0.72 Hz), which rotation rates are much slower than that of PP cell (2 Hz). The parameter values used here are (A) *v*_0_ = 69 [μm/s], *ω_c_* = −6.3 [1/s] and *ω_a_* = 4.6 [1/s], (B) *v*_0_ = 57 [μm/s], *ω_c_* = −4.0 [1/s] and *ω_a_* = 2.1 [1/s], which are obtained from experimental data and Eq. (3). The meanings of blue, red, and black dots and bars in this figure are the same as those in Fig. 4.

Considering the above three results together, it is more likely that the onset of ciliary dominance alteration on the dark side takes place in direct response to a light-off stimulus, rather than as a delayed response to a light-on stimulus.

## DISCUSSION

In this study, we observed initial phototactic turning of *C. reinhardtii* cells after photostimulation with high-speed video recording. As previously shown (Wakabayashi et al., 2011), both PP and NP cells changed the sign of phototaxis when treated with ROS-modulating agents. Our results demonstrate that the sign of phototaxis is determined by which of the two (*cis*- and *trans*-) cilia beats stronger after photoreception, and that the sign reverses depending on whether the dominant cilium begins to beat stronger when the eyespot faces the light source or away from it (Fig. 8). An important factor that modulates when the onset of ciliary dominance occurs is the intracellular amount of ROS. Our mathematical model supports these findings.

**Fig. 8.**
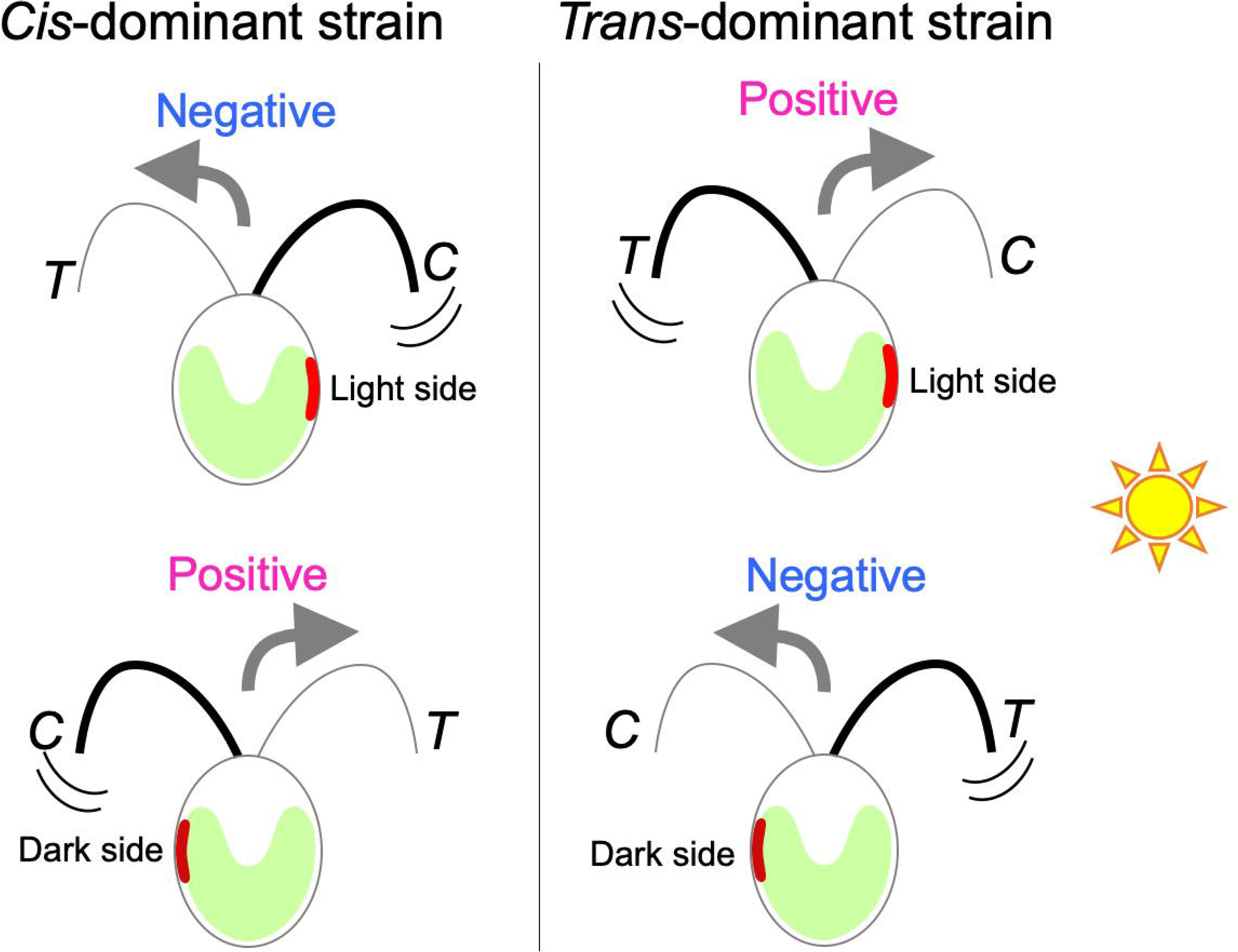
Schematic model of sign-reversal in phototaxis suggested by this study. To make a phototactic turning, *cis*-dominant strain beats the *cis*-cilium (C) stronger than the *trans*-cilium (T), whereas *trans*-dominant strain beats the *trans*-cilium stronger than the *cis*-cilium. The *trans*-dominant strain, such as PP in this study, shows positive phototaxis when the strong beating occurs upon light-on response (i.e. when the eyespot faces the light side) and negative phototaxis when it occurs upon light-off response (i.e. when the eyespot faces the dark side). The *cis*-dominant strain, such as NP in this study, shows negative phototaxis when the strong beating occurs upon light-on response and positive phototaxis when it occurs light-off response.

### The cilium that becomes dominant after photoreception

The first key factor that determines the phototactic sign is whether the *cis*- or the *trans*-cilium is dominant in a given strain. Several previous studies have shown that the two cilia of *C. reinhardtii* are intrinsically different (Kamiya & Hasegawa, 1987; Kamiya & Witman, 1984; Rüffer & Nultsch, 1987, 1991, 1998). Rüffer and Nultsch (1991, 1998) carried out high-speed cinematographic observation on cells trapped with a suction pipette. They found that, upon photo-stimulation, the *trans*-cilium tended to beat more strongly (with a larger amplitude and at a higher frequency) than the other in positively phototactic strains, whereas the *cis*-cilium beat more strongly than the other in negatively phototactic strains (Rüffer & Nultsch, 1991, 1998). Our results, obtained from observation of free-swimming cells undergoing phototactic turning, are basically consistent with these studies.

How is the dominant cilium determined? The results of the present study, as well as those of Rüffer and Nultsch (1991), indicate that the cilium that becomes dominant after photostimulation is the *trans*-cilium in PP and the *cis*-cilium in NP cells. One would be inclined to assume that the Ca^2+^ sensitivities of the *cis-* and *trans*-axonemes (the inner structure of cilia) are reversed between PP and NP. However, in detergent-extracted and motility-reactivated cell models, the Ca^2+^-dependent motility of the two axonemes attached to a single cell body was found not to differ between PP and NP strains (Wakabayashi et al., 2011). Therefore, factors other than axonemes may cause differences in ciliary Ca^2+^-response of PP and NP cells. For example, some detergent-soluble constituents of cilia, or some chemical modification of the axoneme that is not retained after detergent extraction, may be responsible.

Previously, we identified the defect that causes the negatively phototactic phenotype in the NP strain (*agg1*) as the loss of a protein that possibly functions in mitochondria (Ide et al., 2016). If this protein functions in the respiratory chain, the redox poise and/or the amount of ROS could differ between PP and NP cells and differentially modulate the activities of membrane proteins, such as Ca^2+^ channels and pumps. Other *C. reinhardtii* mutants, *agg2* and *agg3*, are also known to display negative phototaxis. Although whether the *cis*- or the *trans*-cilium is dominant in these mutants has not been determined, their defects are also caused by mutations in non-axonemal proteins; the causative protein Agg2 is localized to the proximal ciliary membrane, while Agg3 is a flavodoxin that localizes to the ciliary matrix (Iomini, Li, Mo, Dutcher, & Piperno, 2006). Loss of these proteins may modulate the function of the ciliary membrane and could switch the dominant cilium.

### How is the off-response of the ciliary dominance produced?

The second key factor that determines the phototactic sign is the timing at which the dominant cilium starts to increase power after photoreception. We showed that *t*-BOOH, a membrane-permeable ROS that promotes positive phototaxis, induces the onset of *cis*-cilium dominance on the dark side in NP cells, whereas DMTU, a membrane-permeable ROS-scavenger that promotes negative phototaxis, induces that of the *trans*-cilium dominance on the dark side in PP cells (Fig. 2A, B). The onset of ciliary dominance on the dark side could be interpreted as a so-called off response or a step-down response, which means that cells respond to a light-off stimulus.

Off-response of *C. reinhardtii* has been suggested by Rüffer and Nultsch (1991), but their “off-response” is different from what we observed for the majority of cells showing the sign reversal. They showed that the beating in the two cilia reciprocally changes upon reception of light-on as well as light-off stimuli: i.e., a strain with the *trans* cilium that beats strongly for an on-response will have the *cis* cilium that beats strongly for an off-response. Actually, their result is consistent with a minority of our results. For example, when PP cells showed positive phototaxis, ~30% of the cells beat their cis-cilium strongly when the eyespot faced the dark side (Fig. 2B). How the majority and minority cells switch between the on- and off-responses that Rüffer and Nultsch found was not clear from our experiments.

Off-response itself can be seen in other organisms; for example, in animal’s retina, the cells that normally excite upon light-on response excite upon light-off response when the cells respond to glutamate, a neurotransmitter (Feher, 2012). However, the molecular mechanism of the off response in *C. reinhardtii* is unknown. Theoretically, a response equivalent to an off-response could be accomplished by an appropriate delay in the light-on response. However, our observations on slow-swimming mutants and PP cells in a viscous medium, as well as theoretical considerations, rule out this possibility. Generally, the photoreception by ChR, a light-gated cation channel, is thought to induce depolarization of the cellular membrane. However, the Ca^2+^ influx at the eyespot may induce activation of Ca^2+^-activated K^+^ channel (Vergara et al., 1998), which would induce an increase in K^+^ conductance and concomitant hyperpolarization of the membrane. If the light causes hyperpolarization, a light-off stimulus may induce depolarization of the membrane and elicit an off-response. In addition, recently, ROS-modulating reagents were shown to modulate the phosphorylation state of ChR1 (Bohm et al., 2019). The activity change of ChR1 according to this phosphorylation state may also lead to the off-response. These possibilities can be tested by further electrophysiological analyses using ROS-modulating reagents.

### Mathematical model results support the experimental data

To mathematically assess the plausibility of the Dominant arm model and the Off-response model, we developed a simple model that describes the swimming behaviors of *C. reinhardtii*. The theoretical model and the experimental data both showed that the Off-response model is not accomplished by the presence of a fixed delay time in the light-on response, but by the response to a light-off stimulus (Figs. 5–7). While several previous models have been presented to explain the photobehavior of green alga, ours is one of the simplest, explaining the switching of the phototactic sign through only one equation with some changes of the system parameters (Bennett & Golestanian, 2015; Drescher et al., 2010).

Our model also provides clues to understanding why all the cells of the same strain under the same light conditions do not exhibit the same sign of phototaxis; for example, even when PP cells are treated with H_2_O_2_, which elicits positive phototaxis, ~5% of the cells show negative phototaxis (Wakabayashi et al., 2011). This could be explained by the variance of the delay time in the onset of ciliary dominance. Even though our model precludes the fixed delay time after treatment with the ROS-modulating reagents, the delay time may vary between cells. In Fig. 5, when *γ*_o_ = 1 (i.e., the *trans*-cilium is dominant after photoreception) and *I*_0_ = 0.1 or 0.5, cells show positive phototaxis when ~50 < *τ*_0_ < ~300 msec. The value of τ_0_ has been suggested to be longer than 30~40 msec (Rüffer & Nultsch, 1991; Witman, 1993). Thus, if a cell has longer τ_0_ than 300 msec, which would be caused by several factors including kinetics of Ca^2+^ influx and cellular ROS amounts, this cell will exhibit negative phototaxis. Furthermore, the τ_0_-vz curves in Fig. 4 will change greatly if the eyespot position somewhat varies and causes a change in the τ_0_-vz curve. If a cell has the eyespot at an irregular position, it may exhibit an opposite sign of phototaxis even with the same τ_0_.

We must admit that our mathematical model has some drawbacks: it is based on the major results of the observation. As mentioned earlier, there are exceptional results; for example, ~30% of PP cells showed positive phototaxis with the dominant beating of the *cis* cilium when the eyespot faces the dark side (Fig. 2B). It is still a mystery as to whether cells choose the on- or off-response when exhibiting the same sign of phototaxis. In this study, we took advantage of the fact that ROS-reagents can bias the sign of phototaxis in either direction (Wakabayashi et al., 2011). Similarly, if we can find conditions that bias either the on- or off-reaction, we may be able to derive a model that encompasses all conditions.

In summary, our experimental observations combined with the insights from our theoretical model showed that phototactic signs of *C. reinhardtii* cells are determined by two factors: the genetically determined dominant cilium, and the timing of the onset of strong beating by the dominant cilium after photoreception (Fig. 8). The timing, either on the light side or the dark side, is modulated by the cellular amount of ROS, which is a byproduct of photosynthesis. Cells may monitor photosynthetic activities through ROS amounts, and this regulation mechanism may contribute to maintaining ideal photosynthetic activities by modifying light conditions through phototaxis.

## MATERIALS AND METHODS

### Cell culture and strains

*Chlamydomonas reinhardtii* strains CC-124 (nit1– (nitrate reductase), nit2–, agg1–, mt– (mating type)) (Ide et al., 2016), CC-125 (nit1–, nit2–, mt+), CC-2670 (ida4-, mt+), and CC-2228 (oda1-, mt+) were used. CC-124 and CC-125 were termed PP and NP, respectively. The CC-125 strain maintained in our laboratory appears to have a slight difference in motility characteristics from the same strain available from the Chlamydomonas Resource Center (http://www.chlamycollection.org/) (Sato, Sato, & Toyoshima, 2018; Wakabayashi et al., 2011). CC-2670 (*ida4;* lacking inner-arm dyneins a, c, and d) and CC-2228 (*oda1;* lacking outer-arm dynein and the outer-dynein arm docking complex) were used as slow-swimming mutants. Cells were grown in tris-acetate phosphate medium (TAP) medium with aeration at 25 °C, on a 12 h/12 h light/dark cycle (Gorman & Levine, 1965).

### High-speed observation of phototaxis and measurement of the bodily rotation cycle

Cells were washed with an experimental solution (5 mM Hepes (pH 7.4), 0.2 mM EGTA, 1 mM KCl, 0.3 mM CaCl_2_) (Okita, Isogai, Hirono, Kamiya, & Yoshimura, 2005) and kept under red light for more than 50 minutes before the assays. To induce positive or negative phototaxis, the cell suspensions were treated with tertiary-butylhydroperoxide (*t*-BOOH; final concentration is 0.2 mM) (Wako Pure Chemical Industries) as a ROS reagent, or dimethylthiourea (DMTU; final concentration is 75 mM) (Sigma-Aldrich) as a ROS-scavenging reagent (Wakabayashi et al., 2011). Cell suspensions were put between a coverslip and a glass slide and placed on the stage of a dark-field microscope with an oil-immersion condenser (BX53; Olympus). The directional light to induce phototaxis was produced with two green LEDs (λ=525 nm). The setup is shown in Fig. 2A. First, a weak green light (~5 μmol photons m^-2^ s^-1^) was illuminated (Light 1 in Fig. 2A). Most cells showed either positive or negative phototaxis. Then a stronger green light (Light 2 in Fig. 2A; ~30 μmol photons m^-2^ s^-1^) was illuminated, perpendicular to Light 1. Most of the cells then changed their swimming directions and oriented parallel to the Light 2 beam. The behavior of cells was observed with dim red light (λ >600 nm) and videos were recorded with a high-speed camera (HAS-L2M, DITECT) at 150 fps. The LED for Light 2 was linked with the trigger switch of the high-speed camera so that recording was initiated when it was lit. The timing of photoreception was determined as the time when the eyespot faced the Light 2 side (Fig. 2A, 2B). The position of the eyespot in the helical swimming paths was also determined from the same video footage.

The measurement of the bodily rotation cycle was carried out with the same experimental setup as above (without sideways illuminations). The time required for one bodily rotation was determined from the position of the eyespot on the swimming trajectories, and the rotation period was calculated.

### Phototaxis assay

The phototaxis assay shown in Fig. 5 was carried out by the method described in (Ueki et al., 2016). In brief, cells were washed with the experimental solution and kept under dim red light for 30 min before the phototaxis assays. For dish assays, cell suspensions (~10^7^ cells/mL) were put in Petri dishes (30 mm in diameter, 10 mm thick), illuminated with a green LED (λ=525 nm, ~50 μmol photons m^−2^ s^−1^) from one side for 5 min, and photographed (DSC-RX100M2; Sony). For preparation of viscous media, 4.5% or 9.0% Ficoll (Mw 400; 16006-92, Nacalai) was added to the experimental solution. For single-cell analysis, cells were observed under a dark-field microscope (BX-53, Olympus) under dim red light (λ > 600 nm) and recorded to video using a CCD camera (1129HMN1/3; Wraymer). The angle (θ) between the light direction and the swimming direction was measured for 1.5 s, following illumination with a green LED for 15 s. Images of swimming cells were auto-tracked using Image Hyper software (Science Eye), and angles were measured from the cell trajectories. *t*-BOOH (final concentration of 0.2 mM; Wako Pure Chemical Industries) was used as a ROS reagent, and dimethylthiourea (final concentration of 75 mM; Sigma-Aldrich) was used as a ROS-scavenging reagent

### Electrophysiology

PRCs were assessed in a population of *C. reinhardtii* cells by the method of Sineshchekov et al. (1992) (O. A. Sineshchekov et al., 1992, 1994). In brief, 1 ml of cell suspension in a measuring solution (0.5 mM Hepes, pH 7.3, 0.1 mM CaCl2) was put in a cuvette (10×10×15 mm), with two electrodes on each side of its rectangular bottom. A 500 nm beam of light was generated with an LED source (NSPE510S, Nichia Chemical) and applied from one side of the electrode. The current was measured with a patch-clamp amplifier (Axoclamp 200B, Axon).

### Measurement of ciliary beating frequency

Ciliary beating frequency (CBF) was measured based on the method described in (R Kamiya, 2000) with modifications (Wakabayashi & King, 2006). The median frequency was obtained from the power spectra of fast Fourier-transformed cell body vibration signals in microscopy images averaged for ~20 s.

### Measurement of bodily rotation cycle

Cells were observed under a dark-field microscope with an oil-immersion condenser (BX-53, Olympus) and recorded to video with a high-speed camera (HAS-L2M, DITECT) at 150 fps. The bodily rotation cycle was defined as the time it takes for the eyespot (observed as a bright spot) to return to the same position relative to the cell’s swimming trajectory, and was measured by counting the frames for one cycle.

## Acknowledgments

This work was supported by Japan Society for the Promotion of Science KAKENHI Grants 19H03242, 20K21420, 21H00420, 22H02642, and 22H05674 to KW, 16H06556 to TH, 17H02939 to KS, 17K07370 to KY, 19K23758 to NU, by Ohsumi Frontier Science Foundation to KW, by Global Station for Soft Matter at Hokkaido University to KS and TN, by Dynamic Alliance for Open Innovation Bridging Human, Environment and Materials to TH, KS, TN, and KW and by the Japan Agency for Medical Research and Development (AMED/PRIME) grant JP18gm5810013 to KY.

## References

Bennett, R. R., & Golestanian, R. (2015). A steering mechanism for phototaxis in Chlamydomonas. J R Soc Interface, 12(104), 20141164.

Bohm, M., Boness, D., Fantisch, E., Erhard, H., Frauenholz, J., Kowalzyk, Z., … Kreimer, G. (2019). Channelrhodopsin-1 Phosphorylation Changes with Phototactic Behavior and Responds to Physiological Stimuli in Chlamydomonas. Plant Cell, 31(4), 886–910.

Drescher, K., Goldstein, R. E., & Tuval, I. (2010). Fidelity of adaptive phototaxis. Proc Natl Acad Sci U S A, 107(25), 11171–11176.

Feher, J. (2012). In Quantitative Human Physiology, Elsevier Inc.

Feinleib, M. E. H., & Curry, G. M. (1971). The relationship between stimulus intensity and oriented phototactic response (topotaxis) in Chlamydomonas. In Physiologia Plantarum (Vol. 25, pp. 346–352).

Foster, K. W., & Smyth, R. D. (1980). Light Antennas in phototactic algae. Microbiol Rev, 44(4), 572–630.

Gorman, D. S., & Levine, R. P. (1965). Cytochrome f and plastocyanin: their sequence in the photosynthetic electron transport chain of Chlamydomonas reinhardi. Proc Natl Acad Sci U S A, 54(6), 1665–1669.

Ide, T., Mochiji, S., Ueki, N., Yamaguchi, K., Shigenobu, S., Hirono, M., & Wakabayashi, K.-i. (2016). Identification of the agg1 mutation responsible for negative phototaxis in a “wild-type” strain of Chlamydomonas reinhardtii. Biochemistry and Biophysics Reports, 7, 379–385.

Iomini, C., Li, L., Mo, W., Dutcher, S. K., & Piperno, G. (2006). Two flagellar genes, AGG2 and AGG3, mediate orientation to light in Chlamydomonas. Curr Biol, 16(11), 1147–1153.

Isogai, N., Kamiya, R., & Yoshimura, K. (2000). Dominance between the two flagella during phototactic turning in Chlamydomonas. Zool Sci, 17, 1261–1266.

Kamiya, R. (1988). Mutations at twelve independent loci result in absence of outer dynein arms in Chlamydomonas reinhardtii. J Cell Biol, 107(6 Pt 1), 2253–2258.

Kamiya, R. (2000). Analysis of cell vibration for assessing axonemal motility in Chlamydomonas. Methods, 22, 383–387.

Kamiya, R., & Hasegawa, E. (1987). Intrinsic difference in beat frequency between the two flagella of Chlamydomonas reinhardtii. Exptl. Cell Res., 173, 299–304.

Kamiya, R., Kurimoto, E., & Muto, E. (1991). Two types of Chlamydomonas flagellar mutants missing different components of inner-arm dynein. J Cell Biol, 112(3), 441–447.

Kamiya, R., & Witman, G. B. (1984). Submicromolar levels of calcium control the balance of beating between the two flagella in demembranated models of Chlamydomonas. J Cell Biol, 98(1), 97–107.

Kondo, T., Johnson, C. H., & Hastings, J. W. (1991). Action spectrum for resetting the circadian phototaxis rhythm in the CW15 strain of Chlamydomonas 1. Cells in darkness. Plant Physiol, 95, 197–205.

Landau, L. D., Lifshitz, E. M. (1976). Mechanics, 3rd ed. (Vol. 1 (Course of Theoretical Physics)): Oxford: Butterworth-Heinemann.

Leptpos, K. C., Chioccioli, M., Furlan, S., Pesci, A. I., & Goldstein, R.E. (2018) An Adaptive Flagellar Photoresponse Determines the Dynamics of Accurate Phototactic Steering in Chlamydomonas. bioRxiv, doi: 10.1101/254714 version 2.

Morel-Laurens, N. (1987). Calcium control of phototactic orientation in Chlamydomonas reinhardtii: sign and strength of response. Photochem Photobiol, 45(1), 119–128.

Nagel, G., Ollig, D., Fuhrmann, M., Kateriya, S., Musti, A. M., Bamberg, E., & Hegemann, P. (2002). Channelrhodopsin-1: a light-gated proton channel in green algae. Science, 296(5577), 2395–2398.

Nagel, G., Szellas, T., Huhn, W., Kateriya, S., Adeishvili, N., Berthold, P., … Bamberg, E. (2003). Channelrhodopsin-2, a directly light-gated cation-selective membrane channel. Proc Natl Acad Sci U S A, 100(24), 13940–13945.

Okita, N., Isogai, N., Hirono, M., Kamiya, R., & Yoshimura, K. (2005). Phototactic activity in Chlamydomonas ‘non-phototactic’ mutants deficient in Ca2+-dependent control of flagellar dominance or in inner-arm dynein. J Cell Sci, 118(3), 529–537.

Rüffer, U., & Nultsch, W. (1987). Comparison of the beating of cis-and trans-flagella of Chlamydomonas cells held on micropipettes. Cell Motil Cytoskeleton, 7, 87–93.

Rüffer, U., & Nultsch, W. (1998). Flagellar coordination in Chlamydomonas cells held on micropipettes. Cell Motil Cytoskeleton, 41(4), 297–307.

Rüffer, U., & Nultsch, W. (1991). Flagellar photoresponses of Chlamydomonas cells held on micropipettes: II. Change in flagellar beat pattern. Cell Motility Cytoskeleton, 18(4), 269–278.

Sato, N., Sato, K., & Toyoshima, M. (2018). Analysis and modeling of the inverted bioconvection in Chlamydomonas reinhardtii: emergence of plumes from the layer of accumulated cells. Heliyon, 4(3), e00586.

Sineshchekov, O. A., Govorunova, E. G., Der, A., Keszthelyi, L., & Nultsch, W. (1992). Photoelectric Responses in Phototactic Flagellated Algae Measured in Cell-Suspension. J Photochem Photobiol B-Biology, 13(2), 119–134.

Sineshchekov, O. A., Govorunova, E. G., Der, A., Keszthelyi, L., & Nultsch, W. (1994). Photoinduced electric currents in carotenoid-deficient Chlamydomonas mutants reconstituted with retinal and its analogs. Biophys J, 66(6), 2073–2084.

Sineshchekov, O. A., Jung, K.-H., & Spudich, J. L. (2002). Two rhodopsins mediate phototaxis to low-and high-intensity light in Chlamydomonasreinhardtii. PNAS, 99(13), 8689–8694.

Suzuki, T., Yamasaki, K., Fujita, S., Oda, K., Iseki, M., Yoshida, K., … Takahashi, T. (2003). Archaeal-type rhodopsins in Chlamydomonas: model structure and intracellular localization. Biochem Biophys Res Commun, 301(3), 711–717.

Takada, S., Wilkerson, C. G., Wakabayashi, K., Kamiya, R., & Witman, G. B. (2002). The outer Dynein arm-docking complex: composition and characterization of a subunit (oda1) necessary for outer arm assembly. Mol Biol Cell, 13(3), 1015–1029.

Takahashi, T., & Watanabe, M. (1993). Photosynthesis modulates the sign of phototaxis of wild-type Chlamydomonas reinhardtii. Effects of red background illumination and 3-(3’,4’-dichlorophenyl)-1,1-dimethylurea. FEBS Lett, 336(3), 516–520.

Ueki, N., Ide, T., Mochiji, S., Kobayashi, Y., Tokutsu, R., Ohnishi, N., … Wakabayashi, K. (2016). Eyespot-dependent determination of the phototactic sign in Chlamydomonas reinhardtii. Proc Natl Acad Sci U S A, 113(19), 5299–5304.

Vergara, C., Latorre, R., Marrion, N. V., & Adelman, J. P. (1998). Calcium-activated potassium channels. Curr Opin Neurobiol, 8(3), 321–329.

Wakabayashi, K., & King, S. M. (2006). Modulation of Chlamydomonas reinhardtii flagellar motility by redox poise. J Cell Biol, 173(5), 743–754.

Wakabayashi, K., Misawa, Y., Mochiji, S., & Kamiya, R. (2011). Reduction-oxidation poise regulates the sign of phototaxis in Chlamydomonas reinhardtii. Proc Natl Acad Sci USA, 108(27), 11280–11284.

Witman, G. B. (1993). Chlamydomonas phototaxis. Trends Cell Biol, 3, 403–408.

