## Supplemental text and figures for "Basis for the phototaxis sign reversal in the green alga *Chlamydomonas reinhardtii* studied by high-speed observation"

**Supplementary Information**

**Appendix A: Description of mathematical model of the swimming behaviors of *C. reinhardtii*, focused on the timing of the appearance of ciliary dominance**

We approximate the cell body by a rigid ball, and define the cell body axes as shown in Fig. 3; the vector $\mathbf{c}$ is directed from the center of the cell to the anterior, and the vector $\mathbf{b}$ is perpendicular to $\mathbf{c}$ and directed in the direction of *cis*-cilium within the ciliary beating plane. The vector $\mathbf{a}$ is given by $\mathbf{a} = \mathbf{b} \times \mathbf{c}$, where $\times$ indicates the cross product of vectors, and **a**, **b,** and **c** are all unit vectors. Because these vectors are fixed with the body of *C. reinhardtii* cell, the directions of **a**, **b,** and **c** generally change with time as the cell swims.

Let $\boldsymbol{\omega}(t)$ be the angular velocity of the cell body at time $t$. The time evolutions of **a**, **b,** and **c** are then given as

$$\frac{d\mathbf{a}}{dt} = \boldsymbol{\omega} \times \mathbf{a}$$
$$\frac{d\mathbf{b}}{dt} = \boldsymbol{\omega} \times \mathbf{b}, \quad \text{--[S1]}$$
$$\frac{d\mathbf{c}}{dt} = \boldsymbol{\omega} \times \mathbf{c}$$

The rotational motion of the *C. reinhardtii* cell comes from the beating patterns of two cilia that are fixed at the cell body. Thus it is reasonable to express $\boldsymbol{\omega}(t)$ in terms of the body axes **a**, **b,** and **c** as $\boldsymbol{\omega}(t) = \omega_a(t)\mathbf{a} + \omega_b(t)\mathbf{b} + \omega_c(t)\mathbf{c}$, with some functions of $t$,

$\omega_a(t)$, $\omega_b(t)$ and $\omega_c(t)$. In this model we do not consider the rotation about **b** axis i.e., we put $\omega_b(t) = 0$, because the axis **b** is within the ciliary beating plane and it is expected that the beating of the two cilia does not create the rotation about the axis **b**. In contrast, $\omega_c(t)$ and $\omega_a(t)$ take finete values. In typical forward swimming, *C. reinhardtii* cells rotate about $\mathbf{c}$ with a period, in a counterclockwise direction when viewed from the posterior pole (1). Thus we assume that $\omega_c$ is constant and put $\omega_c(t) = -\omega_c^{(0)}$ [1/s], where $\omega_c^{(0)}$ is a positive constant which is determined from the experimental observation (see Appendix C). In addition to this rotation, another rotational motion about $\mathbf{a}$ can arise, which comes from the difference in beating strengths between the two cilia. When the *trans*-cilium beats more strongly than the *cis*-cilium, a negative component of $\omega_a(t)$ arises, and vice versa. The difference in beating strength of two cilia can depends on the light stimulus received by the eyespot, and $\omega_a$ depends on time in general. Thus we split $\omega_a(t)$ into a constant term $\omega_a^{(0)}$ and time-dependent term $p(t)$, as $\omega_a(t) = \omega_a^{(0)} - p(t)$, where $p(t)$ relates to the photoreception at the eyespot. The constant $\omega_a^{(0)}$ is determined from experimental observation (see Appendix C). The setup described above leads to Eq. 1 in the main text.

According to previous work (2), strains of *C. reinhardtii* that exhibit positive phototaxis under typical conditions (such as PP in this study) have the following properties regarding ciliary beating after photoreception: (i) When the intensity of light received by the eyespot, denoted by $I(t)$, increases with time (step-up light stimulus), the *trans*-cilium beats more strongly than the *cis*-cilium. (ii) When $I(t)$ decreases with time (step-down light stimulus), the *cis*-cilium beats more strongly than the *trans*-cilium. (iii) The response of the ciliary beating occurs after photoreception at the eyespot with some delay (2, 3). This delay time $\tau_0$ is about 30-40 msec under usual conditions; however, as shown in our experiment, $\tau_0$ depends on the cell conditions (Fig. 2B). Among the forms of $p(t)$ that satisfy properties (i)-(iii), we use the following simple form:

$$p(t) = \gamma_0 \frac{dI(t-\tau_0)}{dt}, \text{--[S1.1]}$$

where $\gamma_0$ is a constant that represents the sensitivity to the rate of change of $I(t)$ and $\tau_0$ is a constant that represents the time delay of the response of ciliary beating to the light stimulus. If $\gamma_0$ is positive, the dominant cilium of the cell under step-up stimulus is the *trans*-cilium (as in PP cells), while if $\gamma_0$ is negative, the dominant cilium is the *cis*-cilium (as in NP cells). For simplicity, we assume that $\gamma_0$ takes a value of +1 or -1 only.

In this model the light-off response is represented as follows. In the light-off response the dominant cilium strongly beats when the cell receives the step-down stimulus ( $dI(t)/dt < 0$). Thus if the dominant cilium of the cell is the *trans*-cilium and the cell has the light-off response, this situation is represented by $\gamma_0 = -1$ and a usual value of $\tau_0$ (30-40 msec) in eq. S1.1, while when the dominant cilium is the *cis*-cilium and the system has the light-off response, then this is represented by $\gamma_0 = +1$ and the udual value of $\tau_0$. In the mathematical modeling context, in this way, the setup of the light-off response turns out to be the same as that in the case where the dominant cilium is exchanged. In that sense, this model is useful for investigating the effects of the time delay of response of dominant cilium to photo stimulus on phototaxis.

The relationship between $p(t)$ and $I(t)$ introduced in the work on the adaptive photoresponse in *Volvox* (4) also satisfies properties (i)-(iii) (see SI Appendix D). However, since the time delay of the dominant cilium after the reception of photo stimulus is more explicitly expressed in our model, we use Eq. 2 (eq. S1.1). In our model $I(t)$ is given by

$$I(t) = I_0 \frac{(-\mathbf{e}_{light} \cdot \mathbf{e}_{eyespot} + 1)}{2}, \text{--[S2]}$$

where $I_0$ is a positive constant that represents the maximum light intensity the cell senses, and $\mathbf{e}_{light}$ and $\mathbf{e}_{eyespot}$ are unit vectors that represent the direction (from the center of the cell) of light and the eyespot, respectively. These are given as

$\mathbf{e}_{light} = (0,0,-1)$ and $\mathbf{e}_{eyespot} = (\mathbf{a}+\mathbf{b})/\sqrt{2}$ in this model. Other forms of $I(t)$ such as $I(t) = I_0 \frac{(-\mathbf{e}_{light}\cdot\mathbf{e}_{eyespot}+1)}{2}\Theta(-\mathbf{e}_{light}\cdot\mathbf{e}_{eyespot})$ or $I(t) = I_0 \exp((-\mathbf{e}_{light}\cdot\mathbf{e}_{eyespot}-1)/\alpha_0)$ with the Heaviside function $\Theta(x)$ and positive constant $\alpha_0$ are also possible. However, we have checked that using these forms does not qualitatively change the final results of our simulations. Concerning the translational motion of the cell, we simply set

$$\mathbf{v}(t) = d\mathbf{r}/dt = v_0\mathbf{c}, \text{ ---[S3]}$$

where **v** and **r** are the velocity and position vectors of the center of the cell, respectively. $v_0$ is a positive constant that corresponds to the average velocity of *C. reinhardtii* swimming in the $\mathbf{c}$ direction. The speed $v_0$ is determined from the experimental observation (see Appendix C). The above setup closes the equations in terms of **a**, **b**, **c**, and **r**, and the equations are solved under appropriate initial conditions.

Since we regard the cell body as a rigid body, the directions of body axes **a**, **b**, **c** are described using only three variables (5), for which we use the Euler angles $(\theta_1,\theta_2,\theta_3)$, defined in Fig. 4B, where the $x$, $y$, $z$ axes are the Cartesian coordinate system fixed in space. The explicit x, y, z components of **a**, **b**, **c** in terms of $(\theta_1,\theta_2,\theta_3)$ are given by

$$\begin{aligned}
\mathbf{a} &= (\cos\theta_1\cos\theta_2\cos\theta_3 - \sin\theta_1\sin\theta_3,\ \sin\theta_1\cos\theta_2\cos\theta_3 + \cos\theta_1\sin\theta_3,\ -\sin\theta_2\cos\theta_3)\\
\mathbf{b} &= (-\sin\theta_1\cos\theta_3 - \cos\theta_1\cos\theta_2\sin\theta_3, \cos\theta_1\cos\theta_3 - \sin\theta_1\cos\theta_2\sin\theta_3,\ \sin\theta_2\sin\theta_3) \quad \text{--}\\
\mathbf{c} &= (\cos\theta_1\sin\theta_2,\ \sin\theta_1\sin\theta_2, \cos\theta_2)
\end{aligned}$$

[S3.1]

By using the Euler angles and Eq. 1, Eq. S1 are written as

$$
\begin{aligned}
\dot{\theta}_1 &= -\frac{\cos\theta_3}{\sin\theta_2}(\omega_a^{(0)} - \gamma_0 \frac{dI(t-\tau_0)}{dt}) \\
\dot{\theta}_2 &= \sin\theta_3(\omega_a^{(0)} - \gamma_0 \frac{dI(t-\tau_0)}{dt}) \\
\dot{\theta}_3 &= -\omega_c^{(0)} + \cos\theta_3 \frac{\cos\theta_2}{\sin\theta_2}(\omega_a^{(0)} - \gamma_0 \frac{dI(t-\tau_0)}{dt})
\end{aligned}
\quad ,\text{—[S4]}
$$

where the dot over a variable indicates its time derivative.

### Appendix B: Two steady solutions of Eq. S4 and its linear stability analysis

There are two steady solutions for Eqs. S4 and S3 when $dI/dt = 0$:

$$\begin{aligned} &\theta_1 = -\omega_0 t + \beta_0 \\ &\theta_2 = \arctan(\omega_a^{(0)} / \omega_c^{(0)}) \\ &\theta_3 = 0 \\ &\mathbf{v} = v_0(\frac{\omega_a^{(0)}}{\omega_0}\cos(-\omega_0 t + \beta_0), \frac{\omega_a^{(0)}}{\omega_0}\sin(-\omega_0 t + \beta_0), \frac{\omega_c^{(0)}}{\omega_0}) \end{aligned} \quad \text{,--[S5]}$$

and

$$\begin{aligned} &\theta_1 = \omega_0 t + \beta_0 \\ &\theta_2 = \pi + \arctan(\omega_a^{(0)} / \omega_c^{(0)}) \\ &\theta_3 = 0 \\ &\mathbf{v} = v_0(-\frac{\omega_a^{(0)}}{\omega_0}\cos(\omega_0 t + \beta_0), -\frac{\omega_a^{(0)}}{\omega_0}\sin(\omega_0 t + \beta_0), -\frac{\omega_c^{(0)}}{\omega_0}) \end{aligned} \quad \text{, --[S6]}$$

where

$$\omega_0 = \sqrt{(\omega_c^{(0)})^2 + (\omega_a^{(0)})^2} \quad \text{--[S6.1]}$$

and $\beta_0$ is a constant determined by initial conditions. Solution S5 represents the state where the cell moves in the positive z direction with the constant z velocity component

$$v_z = v_0 \omega_c^{(0)} / \omega_0, \quad \text{——[S6.2]}$$

drawing a left-handed spiral trajectory. This state corresponds to positive phototaxis (note that $\omega_c^{(0)} > 0$ and $\mathbf{e}_{light} = (0, 0, -1)$). Note that in this state $\mathbf{e}_{light} \cdot \mathbf{e}_{eyespot}$ is constant in time, and $I(t)$ does not change with time, which is consistent with the above setup $dI(t)/dt = 0$. Similarly, solution S6 is the state of negative phototaxis, where

$v_z = -v_0\omega_c^{(0)} / \omega_0$. Which state, S5 or S6, appears as a final state (which is the state of the model after a long simulation time) is determined by the values of parameters such as $\gamma_0$, $I_0$, and $\tau_0$. Here, we perform the linear stability analysis of the steady solutions Eqs. S5 and S6 to Eq. S4 which are functional equations for $\theta_1$, $\theta_2$, and $\theta_3$.

First we put $\theta_i = \bar{\theta}_i + \delta\theta_i$ in Eq. S4 with the solution $\bar{\theta}_i$ of Eq. S4 (which $\bar{\theta}_i$ is Eq. S5 or S6) and linearize Eq. S4 with respect to $\delta\theta_i$. We then have

$$\frac{d\delta\theta_i(t)}{dt} = \sum_{j=1}^{3} A_{ij}\delta\theta_j(t) + \sum_{j=1}^{3} B_{ij}\frac{d\delta\theta_j(t-\tau_0)}{dt}, \text{ --[S7]}$$

where $A_{ij}$ and $B_{ij}$ are matrix components that depend on the solution $\bar{\theta}_i$ and the parameters of the system, such as $\gamma_0$, $I_0$, and $\tau_0$. $i$ runs from 1 to 3. Multiplying both sides of Eq. S7 by $e^{-st}$, integrating with $t$ over the range $[0, \infty]$, and rearranging the equations gives

$$\sum_{j=1}^{3}(s\delta_{ij} - A_{ij} - se^{-s\tau_0}B_{ij})\delta\theta_j[s] = \delta\theta_i(0) + \sum_{j=1}^{3} B_{ij}e^{-s\tau_0}(-\delta\theta_j(0) + \int_{-\tau_0}^{0}\delta\theta_j(t)e^{-st}dt), \text{--[S8]}$$

where $\delta\theta_i[s]$ is the Laplace transform of $\delta\theta_i(t)$ given by $\delta\theta_i[s] = \int_0^{\infty}\delta\theta_i(t)e^{-st}dt$, and $\delta\theta_i(t)$ for $-\tau_0 \le t \le 0$ are constants given as initial conditions. Equation S8 indicates that whether $\delta\theta_i(t)$ converges to zero or diverges to infinity when $t$ goes to infinity is determined by roots of the following equation for $s$

$$\det(\mathbf{C}) = 0, \text{ ——[S9]}$$

where $C_{ij} = s\delta_{ij} - A_{ij} - se^{-s\tau_0}B_{ij}$ (6). Equation S9 always has a solution $s = 0$, because the solutions of Eq. S4, Eqs. S5 and S6, are essentially periodic solutions and hence along the periodic orbits any perturbation to the state is neutral. Thus, the roots of Eq. S9 for $s$ other than $s = 0$, which we denote by $s_0$'s, determine the stability of the solution of $\bar{\theta}_i$. If all $s_0$'s have negative real parts, the target solution $\bar{\theta}_i$ is stable, while if even one root $s_0$ has a positive real part, the target solution $\bar{\theta}_i$ is unstable. In Fig. S2 the maximum values of $\mathrm{Re}(s_0)$ for the target solution $\bar{\theta}_i$ for some values of $\gamma_0$ and $I_0$ are plotted as a function of $\tau_0$. The results of the linear stability analysis of the linearized equations S7 agree with those obtained from numerical simulations of the original equations S4 (see Figs 5 and S2). Since $\det(\mathbf{C})$ in eq. S9 is a even function of $\omega_a^{(0)}$, the change in sign of $\omega_a^{(0)}$ does not affect the stability of the steady solution.

**Appendix C: The steady solutions [S5] and [S6] tell us about two issues: (i) determination of the values of $v_0$ , $\omega_a^{(0)}$ and $\omega_c^{(0)}$ in our model determined from the experimental data, (ii) the relative direction of the eyespot of the cell during the steady spiral swimming**

Form eqs. [S5] and [S6] we can get the trajectories of the cell exhibiting the positive and negative phototaxes, respectively, as

$$\mathbf{r}_{positive}(t) = v_0(\frac{\omega_a^{(0)}}{\omega_0^2}\cos(-\omega_0 t + \beta_0 + \frac{\pi}{2}), \frac{\omega_a^{(0)}}{\omega_0^2}\sin(-\omega_0 t + \beta_0 + \frac{\pi}{2}), \frac{\omega_c^{(0)}}{\omega_0} t) \text{——[S10]}$$

$$\mathbf{r}_{negative}(t) = v_0(\frac{\omega_a^{(0)}}{\omega_0^2}\cos(\omega_0 t + \beta_0 + \frac{\pi}{2}), \frac{\omega_a^{(0)}}{\omega_0^2}\sin(\omega_0 t + \beta_0 + \frac{\pi}{2}), -\frac{\omega_c^{(0)}}{\omega_0} t), \text{——[S11]}$$

where we have put the initial positions of the cell to $\mathbf{r}_{positive}(0) = \mathbf{r}_{negative}(0) = v_0 \frac{\omega_a^{(0)}}{\omega_0^2}(\cos(\beta_0 + \frac{\pi}{2}), \sin(\beta_0 + \frac{\pi}{2}), 0)$ . Equations S10 and S11 indicate that the radius $r_0$ of the spiral orbites of the cell during the steady swimming is

$$r_0 = v_0 \frac{|\omega_a^{(0)}|}{\omega_0^2}. \text{——[S12]}$$

In addition,

$$|v_z| = \frac{\omega_c^{(0)}}{\omega_0}. \text{--[S12.1]}$$

The bodily rotation cycle H (Hz) observed in experiments is related to $\omega^{(0)}$ as

$$H = \omega^{(0)} / 2\pi. \text{ –[S13]}$$

Hence if we have the values of $H$, $r_0$ and $v_z$ from the experimental observation, we can determine the values of the model parameters $\omega_a^{(0)}$, $\omega_c^{(0)}$, $v_0$ using eqs. S12, S12.1,

S13, S6.1. The explicit expressions for these quantities are

$$\omega_a^{(0)} = \pm \frac{4\pi^2 H^2 r_0}{\sqrt{{v_z}^2 + 4\pi^2 H^2 {r_0}^2}}$$

$$\omega_c^{(0)} = \frac{2\pi H \,|\, v_z \,|}{\sqrt{{v_z}^2 + 4\pi^2 H^2 {r_0}^2}} \quad . \quad \text{——[S14]}$$

$$v_0 = \sqrt{{v_z}^2 + 4\pi^2 H^2 {r_0}^2}$$

Equation S14 shows that the value of $\omega_a^{(0)}$ is determined only to its absolute value, i.e., the sign of $\omega_a^{(0)}$ is not determined. This result is related to the fact that for some steady helical orbit the cell has the two situations: one is that the eyespot of the cell faces inside the helical orbit and the other is that the eyespot faces outside the helical orbit: both cases are allowed for one helical orbit. This issue is discussed below.

To examine the relation between the direction of the eyespot of the cell and the helical orbit the cell draws in our mathematical we examine the angle between the direction of the eyespot of cell and the radius vector of the helical orbit. First, we investigate the case of positive phototaxis expressed by eq. S10. We project the position vector in eq. S10 onto the x-y plane to have

$$\mathbf{r}_{projection} = v_0 \frac{\omega_a^{(0)}}{{\omega_0}^2}(-\sin\theta_1, \cos\theta_1,\ 0),$$

where $\theta_1 = -\omega_0\, t + \beta_0 + \frac{\pi}{2}$. In a similar way if we project the direction vector of eyespot onto the x-y plane, we have

$$(\mathbf{e}_{eyespot})_{projection} = \frac{1}{\sqrt{2}}(\cos\theta_1 \cos\theta_2 - \sin\theta_1,\ \sin\theta_1 \cos\theta_2 + \cos\theta_1,\ 0),$$

where we have used $\mathbf{e}_{eyespot} = (\mathbf{a}+\mathbf{b})/\sqrt{2}$ and eq. S3.1. From eq. S5, $\theta_2 = \arctan(\omega_a^{(0)} / \omega_c^{(0)})$. Using inner and cross products of $\mathbf{r}_{projection}$ and $(\mathbf{e}_{eyespot})_{projection}$ we can know the angle $\theta_0$ between $\mathbf{r}_{projection}$ and $(\mathbf{e}_{eyespot})_{projection}$ as:

$$\begin{aligned} \cos\theta_0 &= \frac{\mathbf{r}_{projection} \cdot (\mathbf{e}_{eyespot})_{projection}}{|\mathbf{r}_{projection}||(\mathbf{e}_{eyespot})_{projection}|} = \mathrm{sgn}(\omega_a^{(0)}) \frac{\sqrt{(\omega_a^{(0)})^2 + (\omega_c^{(0)})^2}}{\sqrt{(\omega_a^{(0)})^2 + 2(\omega_c^{(0)})^2}} \\ \sin\theta_0 &= \frac{(\mathbf{r}_{projection} \times (\mathbf{e}_{eyespot})_{projection})_z}{|\mathbf{r}_{projection}||(\mathbf{e}_{eyespot})_{projection}|} = -\mathrm{sgn}(\omega_a^{(0)}) \frac{\omega_c^{(0)}}{\sqrt{(\omega_a^{(0)})^2 + 2(\omega_c^{(0)})^2}} \end{aligned} \quad .\text{--[S15]}$$

Equation [S15] indicates that when $\omega_a^{(0)} > 0$ the eyespot faces the outside of the helix orbit, while when $\omega_a^{(0)} < 0$ the eyespot faces the inside of the helix orbit. In other words, depending on the sign of $\omega_a^{(0)}$ the relative direction of the eyespot with respect to the helical orbit changes. The stabilities of the helical orbit for both cases of $\omega_a^{(0)} > 0$ and $\omega_a^{(0)} < 0$ are the same if the absolute value of $\omega_a^{(0)}$ is the same; this is shown in Appendix B. In this way, even when the cell is exhibiting the positive phototaxis, there are two cases where the eyespot faces the inside or outside of the helical orbit. Which situation is chosen is determined by the sign of $\omega_a^{(0)}$.

Next we examine the case of negative phototaxis expressed by eq. S11. In a similar way if we project the position vector of eq. S11 and the direction vector of the eyespot onto the x-y plane, we have

$$\mathbf{r}^{(n)}_{projection} = v_0 \frac{\omega_a^{(0)}}{\omega_0^{\ 2}}(-\sin\theta_1, \cos\theta_1, 0)$$
,
$$(\mathbf{e}_{eyespot})^{(n)}_{projection} = \frac{1}{\sqrt{2}}(\cos\theta_1\cos\theta_2 - \sin\theta_1, \sin\theta_1\cos\theta_2 + \cos\theta_1, 0)$$

where $\theta_2 = \pi + \arctan(\omega_a^{(0)} / \omega_c^{(0)})$. The inner and cross products of these vectors give

$$\cos\theta_0^{(n)} = \frac{\mathbf{r}^{(n)}_{projection}\cdot(\mathbf{e}_{eyespot})^{(n)}_{projection}}{|\mathbf{r}^{(n)}_{projection}||(\mathbf{e}_{eyespot})^{(n)}_{projection}|} = \mathrm{sgn}(\omega_a^{(0)})\frac{\sqrt{(\omega_a^{(0)})^2+(\omega_c^{(0)})^2}}{\sqrt{(\omega_a^{(0)})^2+2(\omega_c^{(0)})^2}}$$
$$\sin\theta_0^{(n)} = \frac{(\mathbf{r}^{(n)}_{projection}\times(\mathbf{e}_{eyespot})^{(n)}_{projection})_z}{|\mathbf{r}^{(n)}_{projection}||(\mathbf{e}_{eyespot})^{(n)}_{projection}|} = \mathrm{sgn}(\omega_a^{(0)})\frac{\omega_c^{(0)}}{\sqrt{(\omega_a^{(0)})^2+2(\omega_c^{(0)})^2}}$$
,--[S16]

where $\theta_0^{(n)}$ is the angle between $\mathbf{r}^{(n)}_{projection}$ and $(\mathbf{e}_{eyespot})^{(n)}_{projection}$. Equation S16 tells us that when $\omega_a^{(0)} > 0$ the eyespot faces the outside of the helical orbit, while when $\omega_a^{(0)} < 0$ the eyespot faces the inside of the helical orbit. As in the case of positive phototaxis, $\det(\mathbf{C})$in eq. S9 for the negative phototaxis expressed by eq. S6 is an even function of $\omega_a^{(0)}$, and hence the change in sign of $\omega_a^{(0)}$ does not change the stability of the steady solution S6. In this way, for both positive and negative phototaxis, there are two cases where the eyespots of the cell faces the inside and outside of the helical orbit, i.e., the direction of the eyespot with respect to the helical orbit reflects the sign of $\omega_a^{(0)}$ of the cell but is not related to the sign of phototaxis.

**Appendix D: Similar properties between the setup, Eq. 2 in our model and the relationship between $p$ and $I$ in the adaptive photoresponse model**

In the adaptive photoresponse model proposed previously (4), the relation between $p(t)$ and $I(t)$ is described by a set of ordinary differential equations:

$$\begin{aligned}\dot{p} &= \frac{1}{\tau_1}(\gamma_1 I - p - h) \\ \dot{h} &= \frac{1}{\tau_2}(\gamma_1 I - h)\end{aligned}, \text{——[S17]}$$

where $\tau_1$, $\tau_2$, and $\gamma_1$ are some positive constants ($\tau_1 > \tau_2$), and $h$ is an additional variable. If we regard $I$ in Eq. S17 as a given function of *t*, Eq. S17 is regarded as a set of nonhomogeneous linear ordinary differential equations for $p$ and $h$. In this situation we can formally solve Eq. S17 as

$$\begin{aligned}p(t) &= p(0)e^{-\frac{t}{\tau_1}} + \frac{\tau_2}{\tau_1 - \tau_2}(e^{-\frac{t}{\tau_2}} - e^{-\frac{t}{\tau_1}})h(0) + \int_0^t B(t-t')I(t')dt' \\ h(t) &= h(0)e^{-\frac{t}{\tau_2}} + \int_0^t \frac{\gamma_1}{\tau_2} e^{-\frac{t-t'}{\tau_2}} I(t')dt'\end{aligned}, \text{--[S18]}$$

where $B(t) = \gamma_1 \frac{\tau_1 e^{-\frac{t}{\tau_2}} - \tau_2 e^{-\frac{t}{\tau_1}}}{\tau_1(\tau_1 - \tau_2)}$ and $p(0)$ and $h(0)$ are initial values of $p$ and $h$, respectively. Integration by parts in $p(t)$ in Eq. S18 gives

$$p(t) = p(0)e^{-\frac{t}{\tau_1}} + \frac{\tau_2}{\tau_1 - \tau_2}(e^{-\frac{t}{\tau_2}} - e^{-\frac{t}{\tau_1}})h(0) + I(0)A(t) + \int_0^t A(t-t')\frac{dI(t')}{dt'}dt', \text{ --[S19]}$$

where

$$A(t) = \frac{\gamma_1\tau_2}{\tau_1 - \tau_2}(e^{-\frac{t}{\tau_1}} - e^{-\frac{t}{\tau_2}}). \text{ –[S20]}$$

The first three terms in Eq. S19 become zeros when $t$ goes to infinity, Thus for $t$ large enough ( $t \gg \tau_1$), $p(t)$ is approximately written as

$$p(t) = \int_{-\infty}^{t} A(t-t')\frac{dI(t')}{dt'}dt'. \text{—[S21]}$$

Since $A$ in Eq. S20 has the properties $A(0) = A(\infty) = 0$, $A(t) > 0$ for $t > 0$, and has a peak at $t = t_0 = \frac{\tau_1\tau_2 \log(\tau_1/\tau_2)}{\tau_1 - \tau_2}$, Eq. S21 indicates that the value of $p$ at time $t$ is mainly contributed by the time derivative of $I$ at time $t - t_0$. In this sense, the relation between $p$ and $I$ in the adaptive response model, Eq. S17, has the time delay with respect to the change in light intensity received by the eyespot, and in that sense, Eq. 2 and Eq. S21 have the same property. Indeed, if we substitute in Eq. S21 $A(t) = \gamma_0\delta(t - \tau_0)$, with some constant $\gamma_0$ and $\tau_0$, where $\delta(t)$ is the delta function, Eq. S21 becomes Eq. 2 in the main text.

### References


1. H. C. Crenshaw, C. N. Ciampaglio, M. McHenry, Analysis of the three-dimensional trajectories of organisms: estimates of velocity, curvature and torsion from positional information. *J Exp Biol* **203**, 961-982 (2000).
2. U. Rüffer, W. Nultsch, Flagellar photoresponses of Chlamydomonas cells held on micropipettes: II. Change in flagellar beat pattern. *Cell motility and the cytoskeleton* **18**, 269-278 (1991).
3. G. B. Witman, Chlamydomonas phototaxis. *Trends Cell Biol* **3**, 403-408 (1993).
4. K. Drescher, R. E. Goldstein, I. Tuval, Fidelity of adaptive phototaxis. *Proc Natl Acad Sci U S A* **107**, 11171-11176 (2010).
5. L. D. Landau, Lifshitz, E. M., *Mechanics, 3rd ed.* (Oxford: Butterworth-Heinemann., 1976), vol. 1 (Course of Theoretical Physics).
6. T. A. Burton, *Stability and periodic solutions of ordinary and functional differential equations* (Orlando, Tokyo, Academic Press, 1985).

### SI Figure legends

**Fig. S1**

**The eyespot position in the helical swimming paths during phototaxis.**

**(A)** Schematic image of the eyespot position in the helical swimming paths. When the *cis*-cilium beats more strongly than the *trans*-cilium, the eyespot points to the outside of the helix (left). When the *trans*-cilium beats more strongly than the *cis*-cilium, the eyespot points to the inside of the helix (right). (B) Proportions of the cells showing positive (P) or negative (N) phototaxis while swimming with the eyespot outside or inside of the helical swimming paths (n=28~62 per condition). Treatment with *t*-BOOH or DMTU did not greatly affect the eyespot position in both strains. (C) Proportions of the PP or NP cells swimming with the eyespot outside or inside of the helical swimming paths (n=30 each) observed under dim red light (i.e. showing no phototaxis).

**Fig. S2**

**Photoreceptor current (PRC) of PP and NP cells with or without ROS-modulating reagents.**

**(A)** Typical data of the photoreceptor current (PRC) measured in PP cell suspension without ROS-modulating reagents. A flashlight was applied at the time indicated by the orange arrow. PRC is determined as the difference between the baseline and the peak value. Delay is determined as the time duration between the flashlight application and the time when the peak current appears. **(B, C)** Effects of treatment with *t*-BOOH (left panels) and DMTU (right panels) on PP **(B)** and NP cells **(C)** on PRC (solid lines) and Delay (dashed lines). The treatment with *t*-BOOH affected neither PRC nor Delay. In contrast, that with DMTU decreased PRC and prolonged Delay.

**Fig. S3**

**The maximum real part of the eigenvalues $s_0$ of the linearized equations of Eq.1 and 2 as a function of $\tau_0$.**

The blue curve is for solution S5, while the red one is for solution S6. If max is negative, the corresponding solution is stable, and vice versa. The results coincide with those given in Fig. 5, where the original Eq. S4 is solved numerically. The parameter values used here are $\omega_c = -11.8$ [1/s] and $\omega_a = 5.8$ [1/s].

**Fig. S4**

The mean velocity $\bar{v}_z$ of the steady-state of the cell as a function of the delay time $\tau_0$ for the two cases: (a) PP in Ficoll400 4.5% and (b) PP in Ficoll400 9.0%. The blue dots exhibit the positive phototaxis, while the red dots exhibit the negative phototaxis. The parameters used here are: (a) , , (b) , , , which are determied by the experimental data in Table 2 by using Eq. (3). The other common parameters are  and .

**Movie S1.**

Movie for Fig. 2A, showing a PP cell making a positive phototactic turning after treatment with *t*-BOOH. The light is illuminated from the right when the green arrow appears. The red circle shows the timing when the eyespot faces the light source side. The movie is run at ×1/10 speed.

**Movie S2.**

Movie for Fig. 2A, showing a NP cell making a positive phototactic turning after treatment with *t*-BOOH. The light is illuminated from the right when the green arrow appears. The red circle shows the timing when the eyespot faces the light source side. The movie is run at ×1/10 speed.

**Movie S3.**

Movie for Fig. 2A, showing a PP cell making a negative phototactic turning after treatment with DMTU. The light is illuminated from the right when the green arrow appears. The red circle shows the timing when the eyespot faces the light source side. The movie is run at ×1/10 speed.

**Movie S4.**

Movie for Fig. 2A, showing a NP cell making a negative phototactic turning after treatment with DMTU. The light is illuminated from the right when the green arrow appears. The red circle shows the timing when the eyespot faces the light source side. The movie is run at ×1/10 speed.

**Movie S5.**

Movie of the numerical simulation of the mathematical model described in SI Appendix A, which shows positive phototaxis when $\tau_0$=0.08 sec. The parameters used here are given in the legend of Fig. 3 (C) in the main text. Space and time scales are indicated in the movie.

**Movie S6.**

Movie of the numerical simulation of the mathematical model described in SI Appendix A, which shows negative phototaxis when $\tau_0$=0.32 sec. The parameters used here are given in the legend of Fig. 3 (D) in the main text. Space and time scales are shown in the movie.

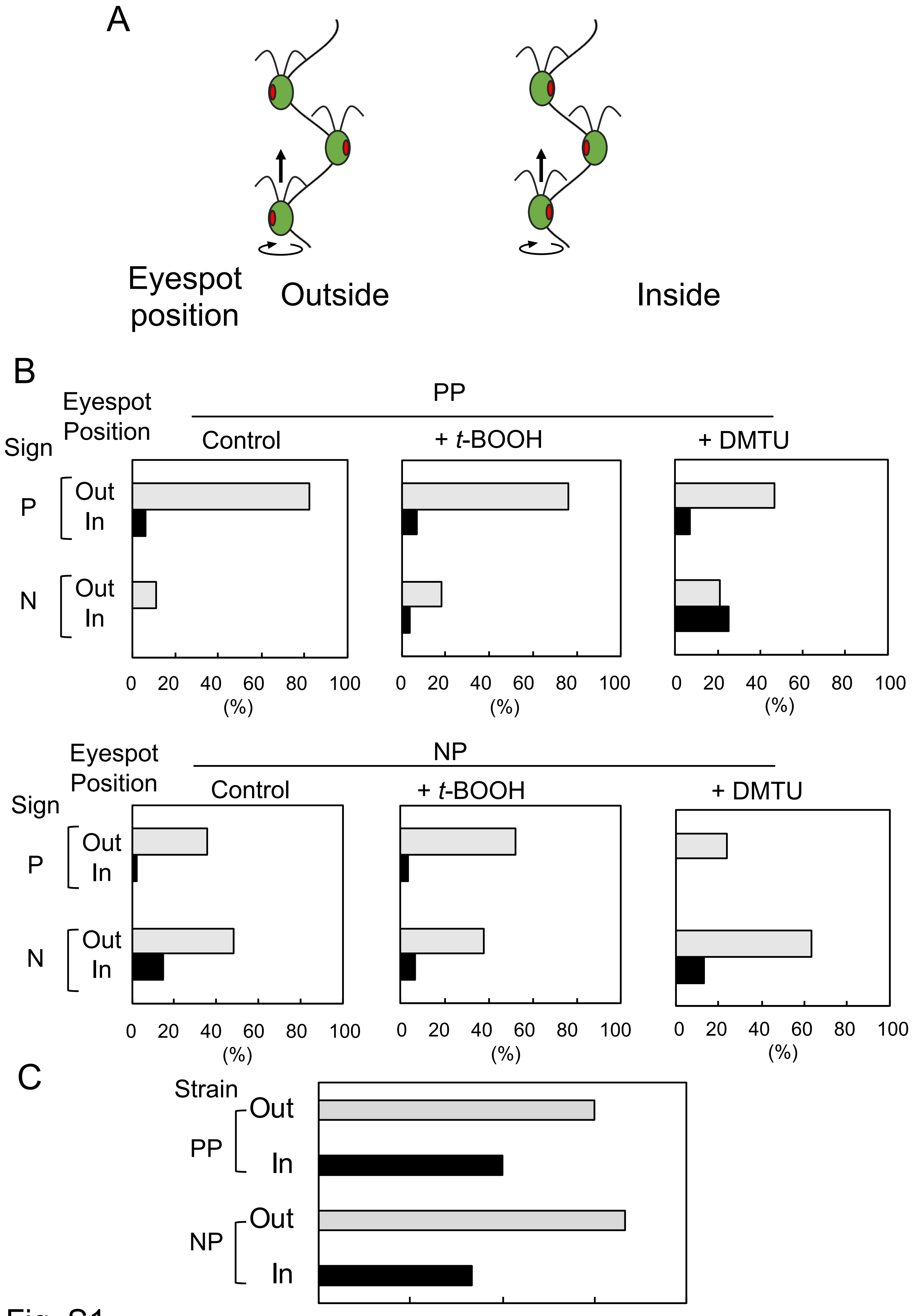
A
Eyespot position
Outside
Inside
B
Eyespot Position
Sign
PP
Control
+ t-BOOH
+ DMTU
P
N
Out
In
0 20 40 60 80 100
(%)
NP
C
Strain
PP
NP
0 20 40 60 80 (%)
Fig. S1

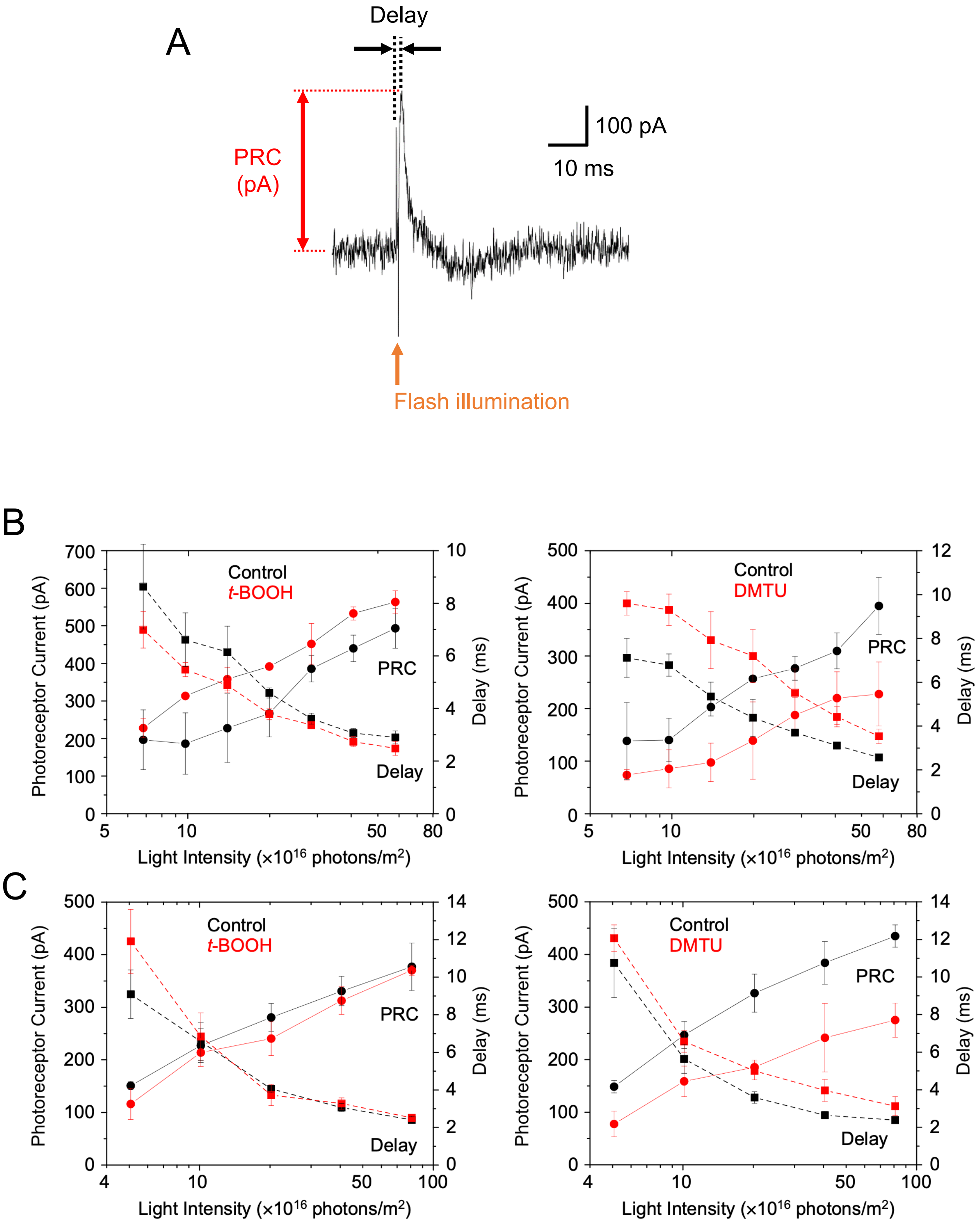
A
Delay
PRC
(pA)
100 pA
10 ms
Flash illumination
B
Control
t-BOOH
PRC
Delay
Photoreceptor Current (pA)
Delay (ms)
Light Intensity (×10^16 photons/m^2)
Control
DMTU
PRC
Delay
Photoreceptor Current (pA)
Delay (ms)
Light Intensity (×10^16 photons/m^2)
C
Control
t-BOOH
PRC
Delay
Photoreceptor Current (pA)
Delay (ms)
Light Intensity (×10^16 photons/m^2)
Control
DMTU
PRC
Delay
Photoreceptor Current (pA)
Delay (ms)
Light Intensity (×10^16 photons/m^2)

Fig. S2

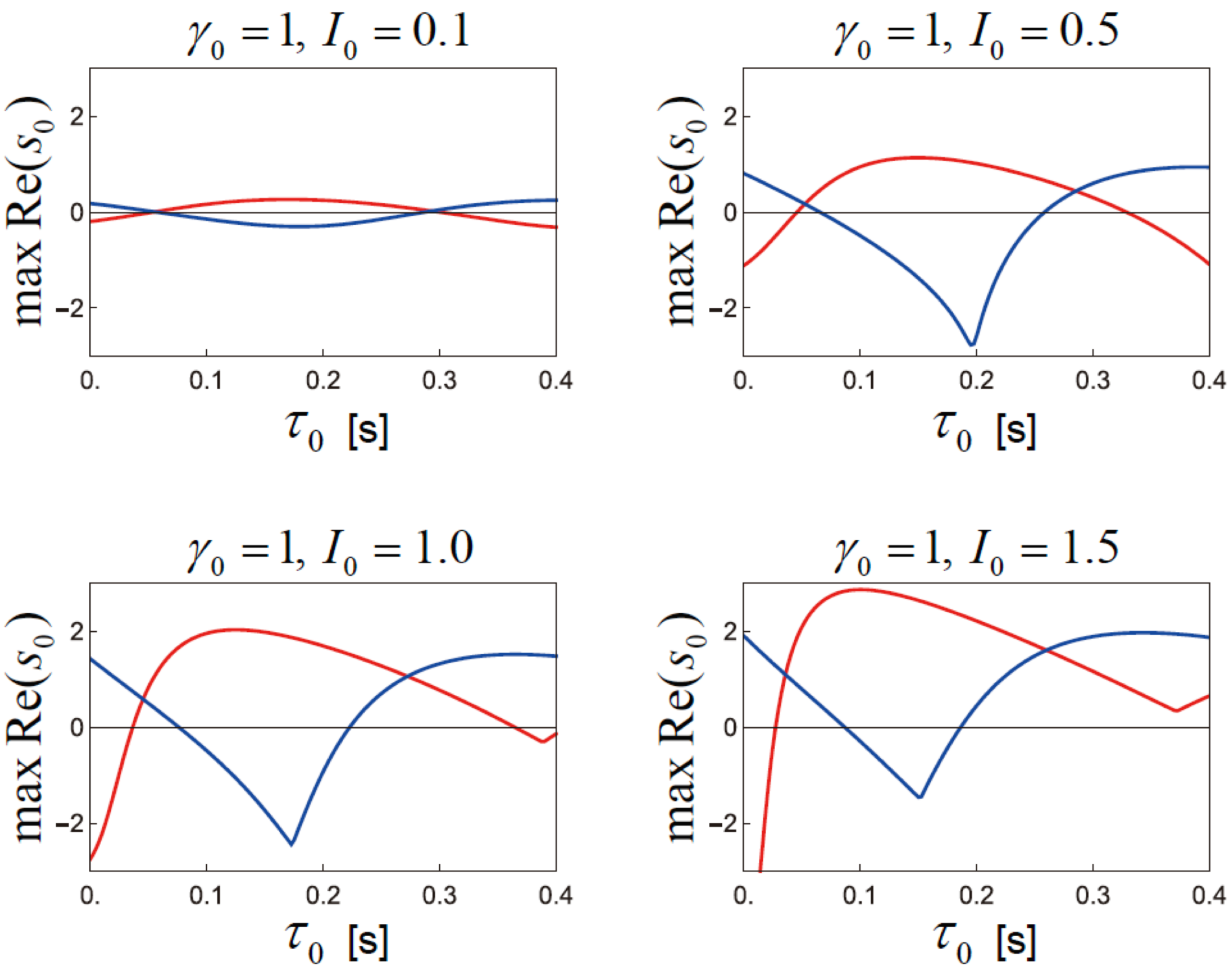
$\gamma_0 = 1,\ I_0 = 0.1$
$\gamma_0 = 1,\ I_0 = 0.5$
$\gamma_0 = 1,\ I_0 = 1.0$
$\gamma_0 = 1,\ I_0 = 1.5$
max Re($s_0$)
2
0
-2
0.
0.1
0.2
0.3
0.4
$\tau_0$ [s]

Fig. S3

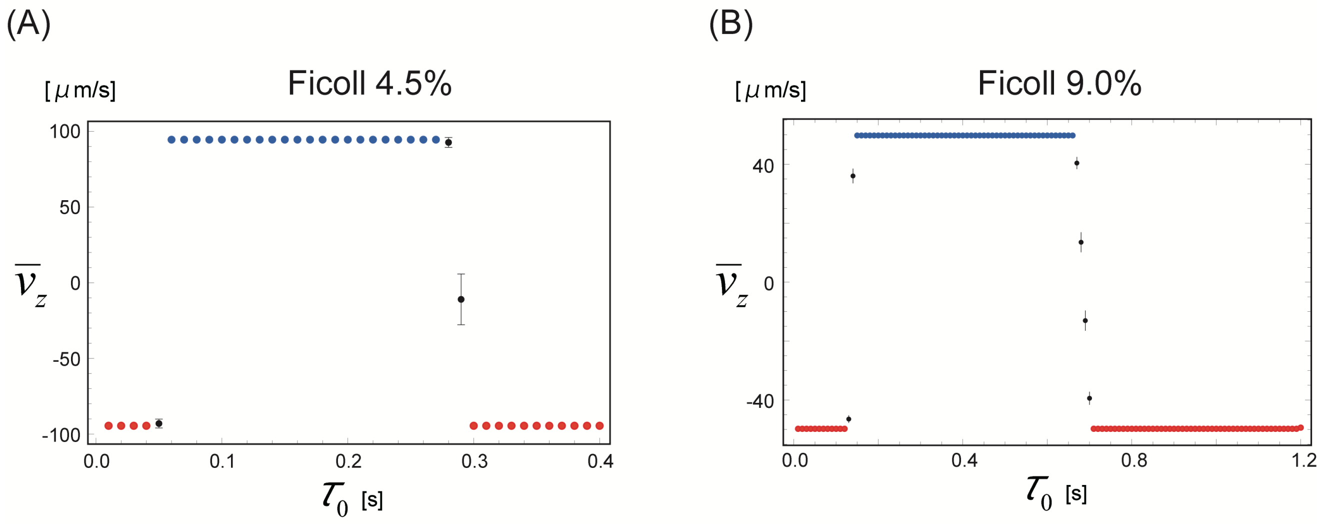

(A)
Ficoll 4.5%
[μm/s]
100
50
0
-50
-100
$\overline{v}_z$
0.0
0.1
0.2
0.3
0.4
$\tau_0$ [s]
(B)
Ficoll 9.0%
[μm/s]
40
0
-40
$\overline{v}_z$
0.0
0.4
0.8
1.2
$\tau_0$ [s]


Fig. S4
